# Developmental Convergence Across Diverse Genetic Architectures in Childhood Apraxia of Speech

**DOI:** 10.64898/2026.10.03.756424

**Authors:** Tiffanie White, Marjan Chapi, Caitlin Raaz, Laurel Bruce, Akshit Modi, Valentin Dinu, Li Liu, Judith Klein-Seetharaman, Beate Peter

**Affiliations:** Krieger School of Arts and Sciences, The Johns Hopkins University, Baltimore, MD, USA; College of Health Solutions, Arizona State University, Phoenix, AZ, USA; Department of Communication Sciences and Disorders, University of Northern Colorado, Greeley, CO, USA

**Keywords:** Childhood Apraxia of Speech, Whole-Exome Sequencing, Early Developmental Brain Transcriptomics, CRISPR interference Perturb-seq, Neurodevelopment, Rare Genetic Variants

## Abstract

Childhood apraxia of speech (CAS) is a neurodevelopmental disorder of speech-motor planning and programming, associated with literacy, academic, and psychosocial difficulties. We performed whole-exome sequencing in 11 families comprising 21 affected and 15 unaffected individuals with phenotyping spanning speech, language, cognition, literacy, and motor function. We identified 60 prioritized variant observations representing 22 unique variants across 20 genes with family-based segregation analysis assessed in informative relatives; 5 arose *de novo*. Our cohort included multiple affected siblings and extended relatives, with several families carrying multiple prioritized variants, suggesting cumulative or oligogenic susceptibility. Speech and literacy impairment co-occurred with oral-motor, feeding, and fine-motor difficulties. BrainSpan co-expression analysis identified *MECP2* and *ANK2* as the genes with the highest observed connectivity in the sampled postnatal network. Exploratory analysis of the KOLF2.1J perturbation atlas in undifferentiated human induced pluripotent stem cells identified a positive *LMO3* expression-response score following *NIPBL* repression and opposing *LMO3* and *ANK2* response directions within a cohesin-associated perturbation cluster. These observations motivate testing in human neural cells. Functional enrichment, protein-association, and developmental-expression analyses supported shared biological themes involving chromatin regulation, neuronal signaling, and extracellular-matrix organization. Among six families, multiple affected relatives shared prioritized variants across a unique set of several genes, demonstrating multilocus candidate profiles whose individual and combined effects may contribute to CAS susceptibility. We propose four novel CAS candidate genes—*LMO3* and *PCDH10*, previously linked to human accelerated regions, as well as *MYCBP2* and *PTPRS*. Our work provides multiple testable hypotheses for future variant-specific functional validation in speech-motor development.

## Introduction

Most children acquire speech naturally without explicit instruction, developing the ability to produce intelligible and coordinated speech sounds by approximately 6 years of age. In contrast, childhood apraxia of speech (CAS) is a rare but severe neurodevelopmental speech sound disorder (SSD) characterized by deficits in the planning and programming of speech movements, resulting in impaired sequencing, timing, and consistency of speech production. CAS is estimated to affect approximately 1 in 1,000 children (Shriberg et al. 2019). CAS is distinguished from other SSDs by its motor planning and coordination deficit rather than impairments in articulatory placement or phonological skills. Affected children often present with inconsistent speech sound errors, disrupted prosody, and difficulty transitioning between sounds and syllables. These deficits frequently extend beyond speech production, contributing to language impairments, literacy challenges, and broader neurodevelopmental difficulties (Miller et al. 2019; Chenausky et al. 2023). These persistent speech deficits frequently require intensive, individualized intervention, although the evidence base for specific CAS treatments remains limited (Morgan et al. 2018).

Early observations dating back to the 1950s noted familial clustering of severe speech disorders, suggesting a heritable component. However, the molecular basis of CAS remained largely unknown until the landmark discovery of pathogenic variants in *FOXP2* in a multigenerational family with severe speech impairment (Lai et al. 2001; Worthey et al. 2013). This discovery established a genetic contribution to CAS and initiated a shift toward molecular characterization of speech and language disorders. The identification of pathogenic *FOXP2* variants established the first monogenic cause of CAS and provided a molecular entry point into the neurobiology of human speech (Takahashi et al. 2009), while the expanding breadth of genetic studies has resulted in overlap with other neurodevelopmental disorders, including epilepsy, intellectual disability (ID), attention-deficit/hyperactivity disorder (ADHD), and autism spectrum disorder (ASD), providing evidence of shared mechanisms across these conditions. Additional genes associated with monogenic forms of CAS continue to be identified through rare-variant analyses. Since the *FOXP2* discovery, advances in genomic sequencing have expanded the number of genes associated with CAS to at least 36 pathogenic variants across 30 genes (Morgan et al. 2024). A subsequent exome study nominated five additional high-confidence candidate genes—*FOXN4, LAMA5, LAMB1, LRRK2,* and *USP17L2*—highlighting substantial genetic heterogeneity (Raaz et al. 2026). In our CAS cohort of 11 probands and their available relatives, the co-occurrence of prioritized variants in multiple biologically relevant genes raises the possibility of multilocus or oligogenic contributions whereby variants in more than one gene may contribute to the observed phenotype. The expanding number of CAS-associated genes demonstrates marked locus heterogeneity in a clinical phenotype with contributions from diverse genes, variant types, inheritance patterns, and potentially multifactorial mechanisms (Kaspi et al. 2023; Morgan et al. 2024). Emerging evidence suggests a critical role for cortico-striatal and cerebellar networks, where disruptions in gene regulation may impair predictive motor control, temporal sequencing, and error correction mechanisms essential for fluent speech production (Todorović et al. 2024).

The cerebellum, in particular, has been increasingly recognized as a key contributor to speech processing. Beyond its primary role in motor coordination, it functions as a predictive system that generates internal models of motor output, compares expected and actual performance, and refines motor commands through error-based learning. Disruption of these processes may contribute directly to the hallmark features of CAS, including impaired timing, sequencing, and speech-motor planning (Peter et al. 2018; Aldinger et al. 2021). In previous phenotypic descriptions of children and adults with CAS, we showed an abundance of error types related to sequential information processing, which is consistent with a cerebellar locus of impairment (Peter et al. 2018). Further supporting a cerebellar locus of impairment, we showed close phenotypic similarities among children with CAS and an adult with a history of cerebellar stroke, in particular deficits in complex motor speech and fine motor tasks and balance (Peter et al. 2021), together supporting the role of the cerebellum in complex motor and communication tasks (Manto et al. 2012; Marien et al. 2014).

Despite marked genetic heterogeneity, genomic studies indicate that genes implicated in CAS converge on neurodevelopmental processes involving chromatin modification, transcriptional regulation, neuronal development, and synaptic function, particularly during early brain development and the formation of neural circuits supporting speech-motor control (Eising et al. 2019; Hildebrand et al. 2020; den Hoed and Fisher 2020; Kaspi et al. 2023). Subsequent reviews have further emphasized the biological overlap among CAS, broader childhood speech disorders, and chromatin-related neurodevelopmental conditions (St John et al. 2023; Morgan et al. 2024). These findings provided the rationale for comparing variants identified in the present cohort with previously reported CAS and neurodevelopmental gene sets. The genes identified in the present cohort and the previously reported CAS gene sets are provided in Supplementary Table S18.

The study by Hildebrand et al. (2020) provided critical evidence that high-confidence variants identified through exome sequencing cluster within co-expressed gene networks active during brain development, supporting a model of convergent transcriptional dysregulation. Similarly, Eising et al. (2019) demonstrated that CAS-associated genes show coordinated expression in the embryonic human brain, further implicating early neurodevelopmental processes in speech acquisition. More recent work has expanded CAS candidate genes and reinforced the importance of shared biological pathways, particularly those related to epigenetic regulation and neuronal development (Kaspi et al. 2023; St John et al. 2023; Formicola et al. 2024). Across previously published high-confidence CAS gene sets, *SETD1A* was reported by Eising et al. (2019) and Kaspi et al. (2023), *SETBP1* by Eising et al. (2019), Hildebrand et al. (2020), and Kaspi et al. (2023), and *DDX3X* by Hildebrand et al. (2020) and Kaspi et al. (2023). These genes illustrate overlap across studies despite substantial genetic heterogeneity. Of these three genes, *SETD1A* was also prioritized in the present cohort, providing a gene-level connection to previous CAS findings.

Across studies, *de novo* variants have been shown to contribute substantially to disease burden, although inherited variants with incomplete penetrance and variable expressivity are increasingly recognized (Morgan et al. 2024). Despite this progress, a significant proportion of cases remain genetically unexplained, highlighting the need for continued gene discovery and functional characterization.

The present study builds upon this established framework by applying comparable sequencing and variant-prioritization strategies to a newly ascertained CAS cohort, enabling both replication of previously implicated genes and identification of novel candidates. Literature-derived CAS gene sets were compiled from Eising et al. (2019), Hildebrand et al. (2020), Kaspi et al. (2023), and related studies for comparative annotation and prioritization; however, variant discovery was not restricted to genes contained in these previously published sets. Variants outside these gene sets remained eligible for prioritization when supported by rarity, predicted functional consequence, inheritance or segregation pattern, and biological relevance to neurodevelopment or speech-language phenotypes. Literature-derived gene sets were used for annotation and comparative interpretation, and prior inclusion in a candidate-gene list was not considered sufficient evidence of pathogenicity.

Given the marked genetic heterogeneity of CAS, interpretation of candidate variants increasingly requires integration of genomic, developmental, and functional evidence rather than reliance on sequence data alone. The present study addresses this gap by examining a clinically characterized cohort of families with CAS using whole-exome sequencing, rigorous variant prioritization, developmental brain gene expression analyses, protein-protein interaction network analysis, CRISPR interference (CRISPRi) perturbations, and gene ontology enrichment. Specifically, we sought to (1) identify and prioritize high-confidence variants with potential relevance to CAS, (2) evaluate both established and novel candidate genes associated with CAS, and (3) compare candidate genes identified in the present CAS cohort with those identified through whole-exome sequencing of three CAS parent– child trios by Raaz et al. (2026), with the overarching goal of determining whether independent genetic findings converge on shared molecular pathways and neurodevelopmental processes relevant to speech and language.

## Materials and Methods

### Ethical approval and consent

Written informed parental permission was obtained for all minor participants, and written informed consent was obtained from all adult participants. Consent included permission for the use of study data in research publications and professional educational activities. This work was conducted with Institutional Review Board oversight at Arizona State University (#STUDY00013638 Biology of Language; #STUDY00010292 Genes, the Brain, and How We Communicate) and the University of Washington (#38237).

### Participants and phenotyping

Twelve independent family units were initially recruited, including 11 CAS families and one additional individual. The final analytic cohort therefore comprised 11 families (21 affected and 15 unaffected individuals). Family sizes in the analytic cohort ranged from 3 to 6 individuals (mean = 3.27). Probands were under 18 years of age at the time of enrollment. CAS diagnosis was established through clinical evaluation by speech-language pathologists, with primary caregiver report indicating persistent speech production deficits as the primary concern.

### Study cohort and family structure

Family structures ranged from trio to multiplex pedigrees, with several families demonstrating multiple affected individuals across generations. One individual was excluded prior to downstream analyses because the participant had a probable diagnosis of Landau-Kleffner syndrome (LKS), a rare epileptic encephalopathy characterized by acquired language regression. Because LKS is a distinct neurological disorder that can independently cause severe speech and language impairment, the participant was excluded to avoid confounding CAS-specific genetic analyses, particularly given their history of epilepsy and prolonged exposure to high-dose antiepileptic medications. No participants or families in the present analytic cohort were included in the cohorts reported by Peter et al. (2016) or Raaz et al. (2026). Although these publications share authors with the present study, they describe distinct participant cohorts. Comparisons with these studies are therefore interpreted as observations across separate cohorts, rather than replication by an independent research team.

### Recorded ancestry and family background

Maternal and paternal ancestry information was obtained from the study’s seqr records. Among the 11 probands, both fields were recorded as “White” for five, “European” for three, and “Hispanic or Latino” for one. One proband had maternal ancestry recorded as “European” and paternal ancestry as “European, Indian”; both fields were unrecorded for the remaining proband. These categories are reproduced as recorded and were not treated as genetically inferred ancestry assignments.

### Phenotyping and behavioral assessment

Behavioral assessment included measures of speech production, language, cognition, literacy, motor coordination, and oral-motor function. Speech articulation was evaluated using the Goldman–Fristoe Test of Articulation–Third Edition (GFTA-3; Goldman and Fristoe 2015), expressive and receptive language using the Comprehensive Assessment of Spoken Language, Second Edition (CASL-2; Carrow-Woolfolk 2017), verbal and nonverbal cognitive abilities using the Reynolds Intellectual Assessment Scales, Second Edition (RIAS-2; Reynolds and Kamphaus 2015), rapid automatized naming using the Rapid Automatized Naming/Rapid Alternating Stimulus Tests (RAN/RAS; Wolf and Denckla 2005), word reading efficiency using the Test of Word Reading Efficiency (TOWRE; Torgesen et al. 2012), fine motor coordination using the Bruininks–Oseretsky Test of Motor Proficiency, Second Edition (BOT-2; Bruininks and Bruininks 2005), and speech-motor performance using alternating motion rate (AMR) and sequential motion rate (SMR) diadochokinetic (DDK) tasks (Fletcher 1972; Fedorenko et al. 2016; Raaz et al. 2026). Complete participant-level linguistic and educational phenotypes are provided in Supplementary Table S1.

Five families consisted of simplex trios, whereas the remaining families demonstrated more complex inheritance patterns. One affected cousin in Family 7 was retained in the pedigree to show family structure but was excluded from the analytic cohort (Fig. 1).

**Fig. 1.**
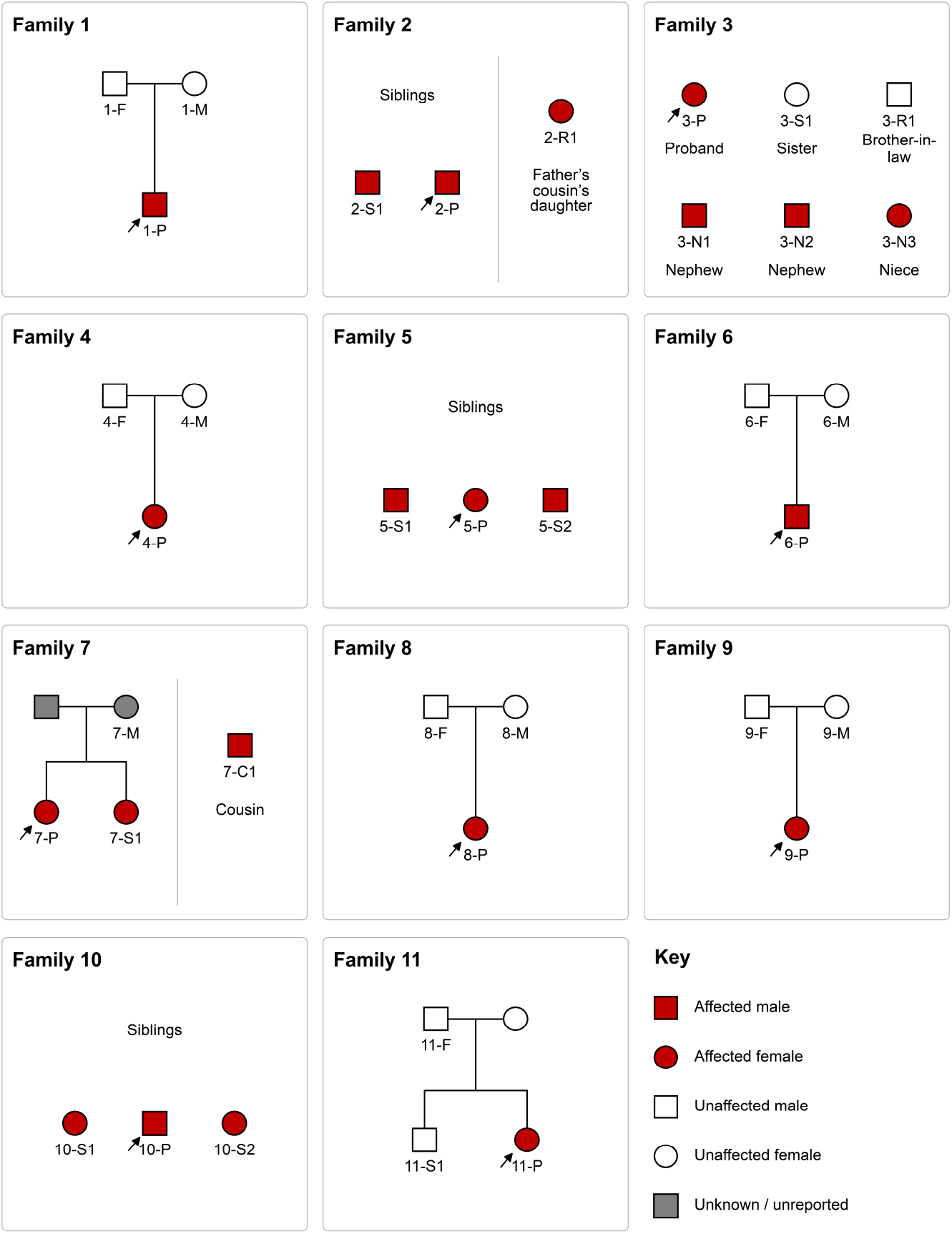
Pedigrees of the 11 families included in the childhood apraxia of speech cohort. Squares denote males and circles denote females; red symbols indicate individuals affected with CAS, white symbols indicate unaffected individuals, and gray symbols indicate unknown status. Arrows identify probands. The cousin in Family 7 (7-C1) was excluded from analysis. P, proband; S, sibling; C, cousin; N, nephew or niece; R, other relative. Unlabeled parents in Family 7 and Mother in Family 11 are shown for family structure. The mother in Family 11 reported no history of speech difficulties and was unaffected phenotypically. Relationship labels refer to the proband; unspecified connecting branches are omitted. Pedigrees follow standardized nomenclature recommendations (Bennett et al. 2022)

### Genetic analyses

Whole-exome sequencing was performed at the University of Washington Center for Rare Disease Research using the Twist Human Exome capture platform (36.5-Mb target) and NovaSeq sequencing. Reads were aligned to GRCh38, and variants were called using Genome Analysis Toolkit (GATK) HaplotypeCaller v4.2.0.0 and annotated using SeattleSeq. Gene symbols were standardized using current symbols approved by the HUGO Gene Nomenclature Committee (HGNC). Variants were filtered in seqr, which is a publicly available platform for identifying candidate disease-causing variants, according to call quality, population frequency, predicted consequence, inheritance, segregation, and phenotypic relevance (Pais et al. 2022; https://seqr.gs.washington.edu). Literature-derived CAS and neurodevelopmental gene sets were used to contextualize and prioritize retained variants but were not used as an exclusionary filter; variants in genes outside these sets were also evaluated when they met the prespecified sequence, inheritance, and biological criteria. Retained variants required read depth >15× and genotype quality >20 and underwent manual review in Integrative Genomics Viewer. Clinical assertions and supporting evidence for retained variants were reviewed in ClinVar (National Center for Biotechnology Information n.d.). Variants in genes with established gene–disease associations were classified according to American College of Medical Genetics (ACMG) and the Association for Molecular Pathology (AMP) guidelines (Richards et al. 2015), whereas variants in genes lacking an established CAS gene–disease association were evaluated using the prespecified evidence framework. Complete sequencing, filtering, prioritization, and classification procedures are provided in Online Resource 1 and summarized in Supplementary Fig. S1.

### Literature-derived comparison gene sets implicated in CAS

A targeted PubMed search was conducted to identify publications relevant to the genetics of CAS, childhood speech disorders, and associated neurodevelopmental conditions. Gene sets used for variant prioritization, comparative analyses, and visualization were derived primarily from the cohort studies of Eising et al. (2019), Hildebrand et al. (2020), and Kaspi et al. (2023), supplemented by the reviews of St John et al. (2023) and Morgan et al. (2024). These sources were selected because they synthesize or report contemporary evidence concerning the genetic architecture and biological pathways associated with CAS and related childhood speech disorders (Supplementary Table S2). Following variant-level filtering and prioritization, a final set of 20 high-confidence candidate genes was retained for downstream comparative and functional analyses. Prior CAS sequencing studies, particularly predominantly trio-based cohorts, identified many high-confidence *de novo* variants, while inherited variants and marked genetic heterogeneity were also reported (Eising et al. 2019; Hildebrand et al. 2020; Kaspi et al. 2023; Morgan et al. 2024). Together with familial aggregation and variable expressivity, these findings raise the possibility that multilocus or oligogenic contributions may operate in a subset of individuals with CAS, although an oligogenic model has not yet been established for the disorder (Peter et al. 2016; Morgan et al. 2024).

Venn and UpSet plots were used to summarize gene-set membership and overlap between the prioritized candidates and published comparison sets (Fig. 2; Supplementary Tables S2 and S18). These comparisons were descriptive and were not tested for enrichment beyond chance. Because the comparison gene sets also informed candidate prioritization, overlap was not interpreted as independent validation of the candidate genes. Evidence concerning shared biological processes was assessed separately through protein-association, functional-enrichment, developmental-expression, and perturbation-response analyses, with each analysis interpreted according to its statistical support and limitations.

**Fig. 2.**
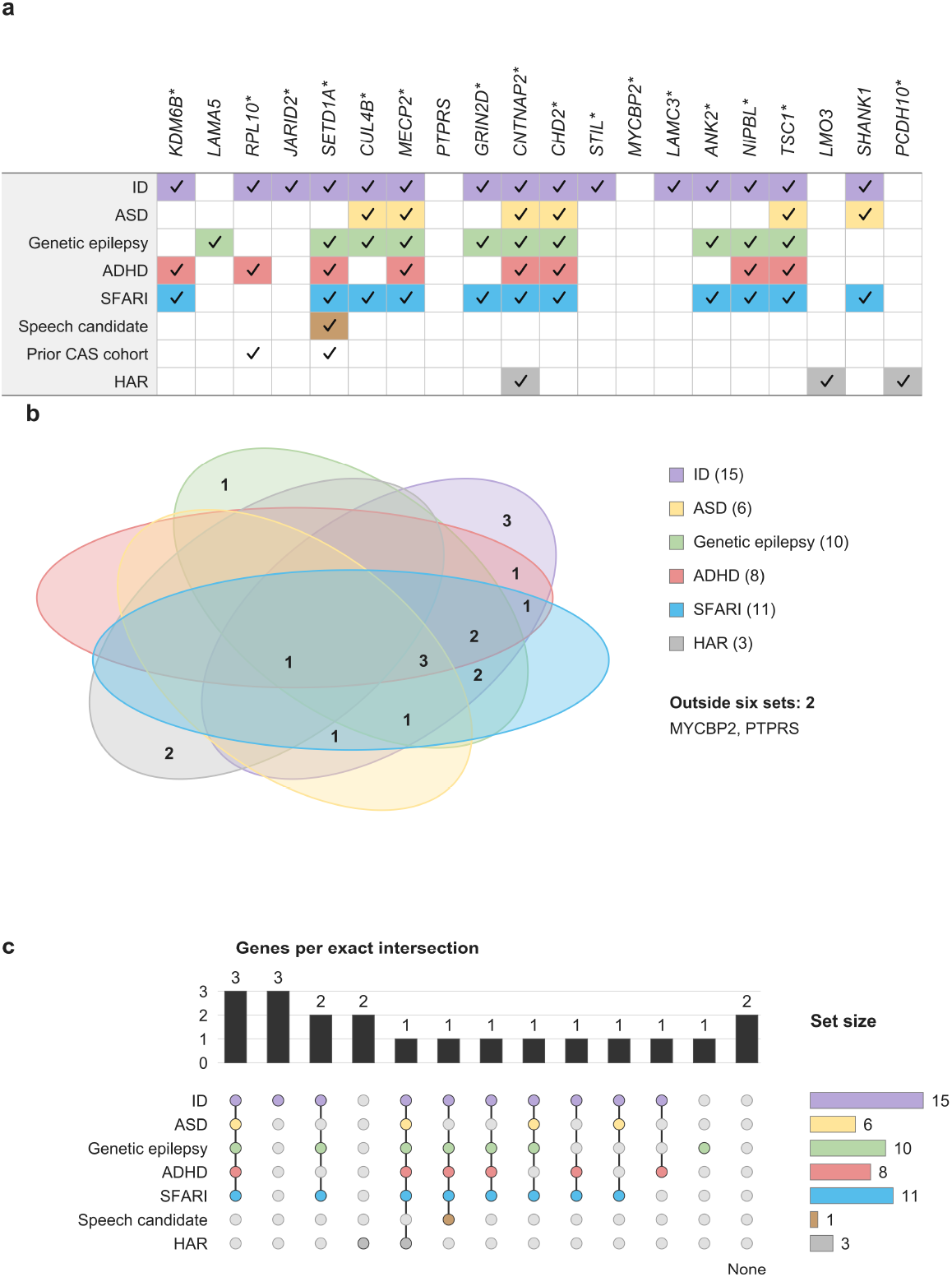
Comparison of CAS candidate genes with published phenotype-associated gene sets. (a) Candidate genes for CAS identified in the present study and by Kaspi et al. (2023); an asterisk identifies genes associated with other neurodevelopmental traits. Each gene is shown once in this table, irrespective of the number of prioritized variants or families in which it was observed. (b) Venn diagram and (c) UpSet plot showing gene-set intersections. The phenotype-associated comparison sets were obtained from Kaspi et al. (2023), with comparison sets and gene-level overlap classifications provided in Supplementary Tables S2 and S18. Intersections describe gene-set membership and were not tested for enrichment beyond chance. The Venn diagram was created using Lucidchart (Lucid Software Inc., South Jordan, UT, USA)

### Collated list of genes of interest

Following variant-level filtering and prioritization, the final high-confidence set comprised 20 candidate genes used for downstream comparative and functional analyses.

High-confidence variants have established evidence supporting a causal relationship with a relevant disorder, whereas some low-confidence variants may be truly pathogenic but currently lack sufficient evidence. Complete low-confidence and predicted damaging missense variant results are provided in Supplementary Tables S3 and S4. Identifying additional cases in future CAS cohorts could strengthen the evidence for these candidate genes and support their reclassification as definitively disease-associated.

### Copy number and structural variants

Using Manta as the calling tool, copy-number and structural-variant data derived from Binary Alignment Map (BAM) and Binary Alignment Map Index (BAI) files were filtered to retain variants marked PASS with cohort allele frequency (AF) < .02. Downstream processing, gene-annotation expansion, exact gene-symbol comparison with the prespecified 20-gene high-confidence CAS candidate set, and record summarization were performed in R version 4.6.0 using *dplyr*, *tidyr*, and *readr*. Candidate copy-number variants (CNVs) were reviewed manually in Integrative Genomics Viewer (IGV). This analysis was included because copy-number variants have been implicated in CAS and other speech sound disorders (Chan et al. 2024).

### KOLF2.1J Perturb-seq Cell Atlas

The CRISPRi Perturb-seq dataset for KOLF2.1J hiPSCs is publicly available via the KOLF2.1J Perturbation Cell Atlas https://y-doctor.github.io/KOLF2.1J_Perturbation_Cell_Atlas/. The KOLF2.1J Perturbation Cell Atlas (Nourreddine et al. 2026) was used as an orthogonal functional screen to assess regulatory connectivity between high-confidence CAS candidates and genes reported in previous CAS and neurodevelopmental studies. This large-scale CRISPRi Perturb-seq resource includes more than 10,000 perturbed genes and 38,000 measured expression targets in human induced pluripotent stem cells. Although human induced pluripotent stem cells (hiPSCs) represent an early pluripotent state rather than a defined fetal developmental stage, the atlas enables systematic assessment of transcriptional responses following gene repression. All 20 high-confidence candidate genes were represented as expression targets, and 13 were directly perturbed (Supplementary Table S19). Fig. 6c presents selected atlas-derived transcriptional-response associations, with no more than two downstream targets displayed per perturbed CAS candidate to maintain readability. Scores and rankings were interpreted descriptively rather than as statistical-significance criteria. Gene-set definitions, relationship inclusion criteria, and visualization procedures are detailed in Online Resource 1 and Supplementary Tables S19–S22.

### Functional data analysis

The BrainSpan Atlas of the Developing Human Brain is publicly available through the Allen Institute for Brain Science at https://www.brainspan.org/rnaseq/search/index.html. Protein–protein interaction analysis was performed using STRING (Search Tool for the Retrieval of Interacting Genes/Proteins; https://string-db.org/) with Homo sapiens as the reference organism and no additional interactors. The full STRING network incorporated both physical and functional associations from curated databases, experimental evidence, co-expression, genomic-context predictions, and text mining.

### Statistical analysis

No statistical methods were used to predetermine the cohort sample size. Randomization was not applicable because participants were grouped according to clinical and genetic characteristics. The BrainSpan and KOLF2.1J analyses used publicly available datasets. Data distributions were not assumed to be normal. Statistical analyses and figure generation were performed in R version 4.6.0 (R Foundation for Statistical Computing, Vienna, Austria). All statistical tests were two-sided, and statistical significance was defined as *p* < .05 unless otherwise specified. Network edges were defined as absolute Spearman correlation coefficients of |ρ| ≥ .50. Complete supplementary methods and results and Supplementary Fig. S1 are provided in Online Resource 1. Supplementary Tables S1–S27 are provided in Online Resource 2.

## Results

### Phenotyping of CAS patients

Behavioral testing demonstrated deficits consistent with a neurodevelopmental speech disorder. The majority of participants exhibited oral-motor impairment and a history of feeding difficulties, and all were diagnosed with CAS by a speech-language pathologist. Speech articulation scores ranged from severely impaired to average on the Goldman–Fristoe Test of Articulation–Third Edition (GFTA-3; Goldman and Fristoe 2015), with standard scores ranging from 40 to 107, while literacy measures on the Test of Word Reading Efficiency (TOWRE; Torgesen et al. 2012) similarly ranged from impaired to average (standard scores: 67–107). Verbal and nonverbal cognitive abilities, assessed using the Reynolds Intellectual Assessment Scales, Second Edition (RIAS-2; Reynolds and Kamphaus 2015), generally ranged from low average to average, although individual scores extended beyond these ranges (Verbal Intelligence Index, 75–112; Nonverbal Intelligence Index, 66–123), indicating that speech and literacy impairments occurred despite largely preserved cognitive functioning. Fine motor coordination was assessed using the Manual Coordination composite of the Bruininks–Oseretsky Test of Motor Proficiency, Second Edition (BOT-2; Bruininks and Bruininks 2005). Scores ranged from 39 to 61 (normative mean = 50, SD = 10), with most scores falling below the normative mean. Oral and speech-motor findings further supported a motor-planning phenotype. Recurrent abnormalities included difficulty with rapid alternating oral movements, reduced jaw–tongue dissociation during tongue lateralization, impaired lip protrusion or alternating smile–pucker movements, and difficulty coordinating smiling and swallowing. Diadochokinetic testing showed poorer sequential (disyllabic) than alternating (monosyllabic) repetition rates in most participants, consistent with impaired sequential motor processing previously described as a potential CAS-related endophenotype by Peter and colleagues. Feeding difficulties were also frequently reported, suggesting that motor involvement extended beyond speech production in a subset of affected individuals. Together, these findings support a broader motor phenotype involving speech, oral-motor, and fine-motor systems rather than an isolated articulation deficit. This pattern is consistent with previous work demonstrating that CAS can involve motor discoordination across speech, fine-motor, and gross-motor domains (Peter et al. 2013; Iuzzini-Seigel 2019; Peter et al. 2021; Raaz et al. 2026).

### Genotype–phenotype relationships among families with CAS

CAS represented the shared core phenotype across affected individuals, whereas associated neurodevelopmental features showed variable expression within and between families. Gross- and fine-motor delays were present in subsets of affected individuals, while no vision impairment, hearing loss, or seizures were reported. Recurrent oral-motor abnormalities—including reduced jaw–tongue dissociation, difficulty sequencing alternating oral movements, and impaired smile–swallow coordination—suggested a shared motor-planning phenotype despite substantial genetic heterogeneity. This phenotypic pattern is consistent with previous studies of familial CAS in which deficits in motor sequencing extended across speech and nonspeech tasks and were investigated as potential endophenotypes of genetically mediated speech disorders (Peter et al. 2013; Peter et al. 2016).

A notable gene-level overlap with previous familial CAS research involved *NIPBL*. Peter et al. (2016) identified a segregating *NIPBL* variant in a multigenerational family with CAS, although it was interpreted as one of several variants that might contribute additively to the phenotype rather than as an independently established causal variant. In Family 10, all three affected relatives—two females and one male—shared prioritized variants in *MYCBP2, LAMC3, PTPRS, TSC1, RPL10,* and *NIPBL*, including *NIPBL* c.535G>A (p.Ala179Thr), together with two *ANK2* missense variants: c.4315G>T (p.Gly1439Cys) and c.9900C>A (p.Ser3300Arg). These genes converge across complementary biological layers. *NIPBL* encodes a component of the *NIPBL–MAU2* cohesin-loading complex, which facilitates cohesin loading onto DNA (Parenti et al. 2020). Cohesin is a ring-shaped protein complex that holds sister chromatids together after DNA replication, helps organize chromosomes into loops, and supports DNA repair. Because cohesin does not efficiently bind chromosomes on its own, *NIPBL* assists with the process. This regulates three-dimensional chromatin organization, whereas *ANK2* encodes a neuronal membrane–cytoskeletal scaffold. *LAMC3*, an extracellular-matrix laminin gene, is implicated in cortical organization, while *PTPRS* encodes a receptor-type protein tyrosine phosphatase involved in cell adhesion and neural signaling.

This shared multivariant profile is compatible with a cumulative or multilocus model, although the relative phenotypic contribution of each variant remains unresolved. The independent identification of *NIPBL* in two families affected by CAS provides gene-level recurrence and strengthens its candidacy for involvement in speech and language neurodevelopment. Nevertheless, because multiple variants segregated in both studies, recurrence alone does not establish *NIPBL* as a monogenic cause of CAS (Peter et al. 2016). The presence of multiple segregating variants in both families favors cautious interpretation within a potentially oligogenic model.

### Summary of major neurodevelopmental phenotypes observed across affected individuals

CAS was present in all individuals, whereas gross motor delay and fine motor delay showed variable penetrance. No vision impairment, hearing loss, or seizures were reported. Recurrent oral-motor abnormalities were observed across families, including impaired jaw–tongue dissociation, difficulty with alternating oral movements, and impaired smile–swallow coordination, suggesting a shared motor-planning endophenotype associated with CAS. Segregating candidate variants differed among families, with multiple prioritized variants observed in Families 2, 3, 5, 7, 10, and 11. In Family 3, the *CUL4B* c.140G>C variant was maternally transmitted to the two affected male nephews through their sequenced mother, while their father did not carry the variant. In Family 7, the *STIL* c.340C>T variant was maternally inherited and shared by both affected sisters, whereas the *CHD2* c.239C>T variant was shared by both affected sisters but was absent from their sequenced mother. Across the broader set of families analyzed contemporaneously in seqr, *LAMA5* was recurrent, with one finding in Family 2 of the present cohort and another in a separately reported family (Raaz et al. 2026). This comparison involves distinct families analyzed within a shared research and analysis context; the recurrence of *LAMA5* is therefore presented as a cross-cohort observation rather than an independent analytical replication.

Recurrent findings included *RPL10* in Families 2, 3, 5, 6, and 10 and *PTPRS* in Families 5 and 10. Family 10 was particularly notable because three affected relatives shared variants involving *LAMC3, NIPBL, ANK2,* and other neurodevelopmentally relevant candidates (Peter et al. 2016). Thus, a relatively consistent CAS and motor-planning phenotype occurred in the context of considerable locus and allelic heterogeneity. This combination of phenotypic convergence and genetic heterogeneity parallels recent observations by Raaz et al. (2026), who described three unrelated individuals with CAS and accompanying fine- and gross-motor involvement despite different candidate genetic variants (Raaz et al. 2026; Supplementary Table S1).

### Identification of high-confidence CAS gene variants

Exome sequencing identified high-confidence variants in 9 of the 11 (81.8%) CAS probands, comprising 60 variant observations across the cohort, reflecting the 5 *de novo* and 55 familial variants, indicating contributions from both sporadic mutations and inherited susceptibility to CAS. These observations represented 22 unique variants: 3 stop-gained, 1 in-frame deletion, 1 frameshift, 1 splice-acceptor, and 16 missense variants. Of the total high-confidence variants, 5 arose *de novo*: *KDM6B, LMO3, MECP2, SHANK1,* and *PCDH10,* while 55 were inherited. The *de novo MECP2* c.1393C>T (p.Arg465Ter) variant introduces a premature stop codon and removes a CpG dinucleotide through a CGA-to-TGA substitution, a loss of a potential methylation site through the sequence change with potential downstream effects on methylation-dependent gene regulation. Its local methylation consequence was not assessed in the present study, and the principal predicted consequence remains truncation of the *MECP2* coding sequence.

In silico evidence supported the potential functional significance of these variants, with 77.3% having CADD scores ≥20 and 31.8% having scores ≥30. At the gene level, 80% of prioritized genes had pLI scores ≥.90, and 65% had LOEUF values <.35, indicating substantial intolerance to genetic variation. Together, these findings support the biological plausibility of the prioritized candidates and are consistent with patterns observed among genes implicated in neurodevelopmental disorders. *LMO3* p.Arg167Cys met an ultra-rare allele-frequency threshold of <.0001 in both the exome and genome datasets, and *SHANK1* p.Ala6Thr was rare, with frequencies of .000199 and .0000986, respectively, as reported by VarSome (Kopanos et al. 2019; https://varsome.com).

The identified variants involved established CAS-associated genes (e.g., *KDM6B, JARID2, SETD1A, CUL4B, CHD2, ANK2, NIPBL,* and *RPL10*), neurodevelopmental disorder genes with reported or potential speech-language involvement (e.g., *MECP2, GRIN2D, CNTNAP2, LAMA5, LAMC3, SHANK1, STIL,* and *TSC1*), and the human accelerated region (HAR) associated CAS candidates *LMO3* and *PCDH10*. HARs are evolutionarily conserved genomic elements that acquired rapid sequence changes in humans and often regulate genes involved in brain development and cognition. The novel CAS candidates *PTPRS* and *MYCBP2*, not previously implicated in CAS to our knowledge, further expand the candidate-gene landscape. Key contributing genes included *SETD1A, MECP2,* and *CHD2,* which are associated with chromatin and transcriptional regulation, and *SHANK1* and *GRIN2D,* which are associated with synaptic signaling, supporting convergence on neurodevelopmental and synaptic pathways.

### Prioritization of genes in prior studies

To prioritize variants with potential relevance to CAS, candidate genes were compared against a previously curated panel of 2,147 genes associated with CAS and related neurodevelopmental disorders (Supplementary Table S2). This panel integrates evidence from published CAS sequencing studies, genes implicated in individuals with CAS, genes associated with intellectual disability, epilepsy, and cleft palate, and autism-associated genes cataloged in the Simons Foundation Autism Research Initiative (SFARI) database (Kaspi et al. 2023). After identifying candidate variants in the present study, they were first evaluated with reference to this gene set.

### Copy number and structural variants

CNV analysis identified 362 filtered per-sample structural-variant records across 36 sequenced samples, comprising 142 deletions, 21 duplications, 198 breakend records, and one insertion. These records corresponded to 222 distinct CNV identifiers, and 299 records contained at least one gene annotation. No annotated CNV overlapped the prespecified 20-gene high-confidence CAS candidate set by exact gene-symbol matching. Manual review in IGV identified no CNVs considered plausibly relevant to the observed phenotype.

### Recurrent observations in *RPL10* and *PTPRS*

Analysis of recurrent variants identified *RPL10* as the most frequently observed candidate gene in the cohort. The X-linked *RPL10* p.Arg210Trp variant was identified in five unrelated families and in 12 affected individuals across both sexes, making it the predominant recurrent variant detected. In contrast, the novel candidate gene *PTPRS* harbored two independent heterozygous missense variants (p.Thr1510Met and p.Ala591Thr) identified in two separate families, each segregating with affected relatives in a pattern consistent with autosomal-dominant inheritance. Although the genes harboring recurrent variants have distinct biological functions, they both contribute to fundamental neurodevelopmental processes. *RPL10* encodes a component of the large ribosomal subunit involved in protein synthesis, whereas *PTPRS* encodes a receptor protein tyrosine phosphatase that regulates axon guidance, synaptic organization, and neuronal connectivity. This variant was uncommon in TOPMed, with an allele frequency of 0.000328686 (approximately 0.03287%) recorded in the original variant annotation. Recorded ancestry categories were broad and did not establish genetic independence among families. Shared ancestry or population structure therefore remains a possible contributor to the recurrence of RPL10 p.Arg210Trp across families. Together with two distinct *PTPRS* missense variants prioritized under the study framework, these observations highlight candidate contributions of translational and synaptic pathways to the genetic architecture of CAS.

## Differentially expressed genes

BrainSpan source expression data, gene-expression summaries, regional mappings, sample metadata, developmental comparisons, and network statistics are provided in Supplementary Tables S5–S17.

BrainSpan co-expression analysis combined the 20 research-prioritized CAS candidate genes from the present study with previously reported CAS candidate genes, yielding 53 unique genes after removal of duplicates (Eising et al. 2019; Hildebrand et al. 2020; Kaspi et al. 2023; Supplementary Table S18). Across the 1,378 unique gene pairs in this combined set, median absolute Spearman correlation was 0.2882 in the prenatal network and 0.3381 in the postnatal network, an observed difference of 0.0499 (Fig. 3a, b; Supplementary Table S16). A donor-cluster bootstrap with 10,000 replicates yielded an approximate 95% confidence interval of −0.0599 to 0.1168. Sensitivity analyses restricted to brain regions represented in both developmental groups and adjusted for regional mean ranks also produced intervals that included zero. Descriptively, the proportion of positive correlations increased from 69.0% in the prenatal network to 79.3% in the postnatal network, while the number of gene pairs meeting the network inclusion threshold of |ρ| ≥ 0.50 increased from 328 of 1,378 pairs (23.8%) to 387 of 1,378 pairs (28.1%). In the postnatal network, *MECP2* and *ANK2* had the highest observed connectivity, with 29 edges each, followed by *TRIO* with 28 and *DIP2C* with 26, while *TSC1* had 21 edges. Novel CAS candidate genes *MYCBP2* and *PTPRS* had 26 and 20 edges, respectively. *PTPRS* exceeded both the network mean connectivity of 14.6 and the median connectivity of 15 and was tied at rank 12 among the 53 compared genes, whereas *MYCBP2* ranked fourth (Supplementary Table S17). These connectivity values place both *MYCBP2* and *PTPRS* within a broadly connected developmental transcriptional network but do not establish direct molecular interactions, pathogenicity, or a specific role in CAS. A separate sample-count sensitivity analysis showed that the higher median correlation in the sampled postnatal network persisted when the number of postnatal tissue samples was reduced to match the prenatal sample count (Online Resource 1). However, this analysis did not address repeated sampling within donors or dependence among gene-pair correlations. Accordingly, the sampled networks differed descriptively, but the donor-aware analyses did not provide clear evidence of a population-level developmental increase.

**Fig. 3.**
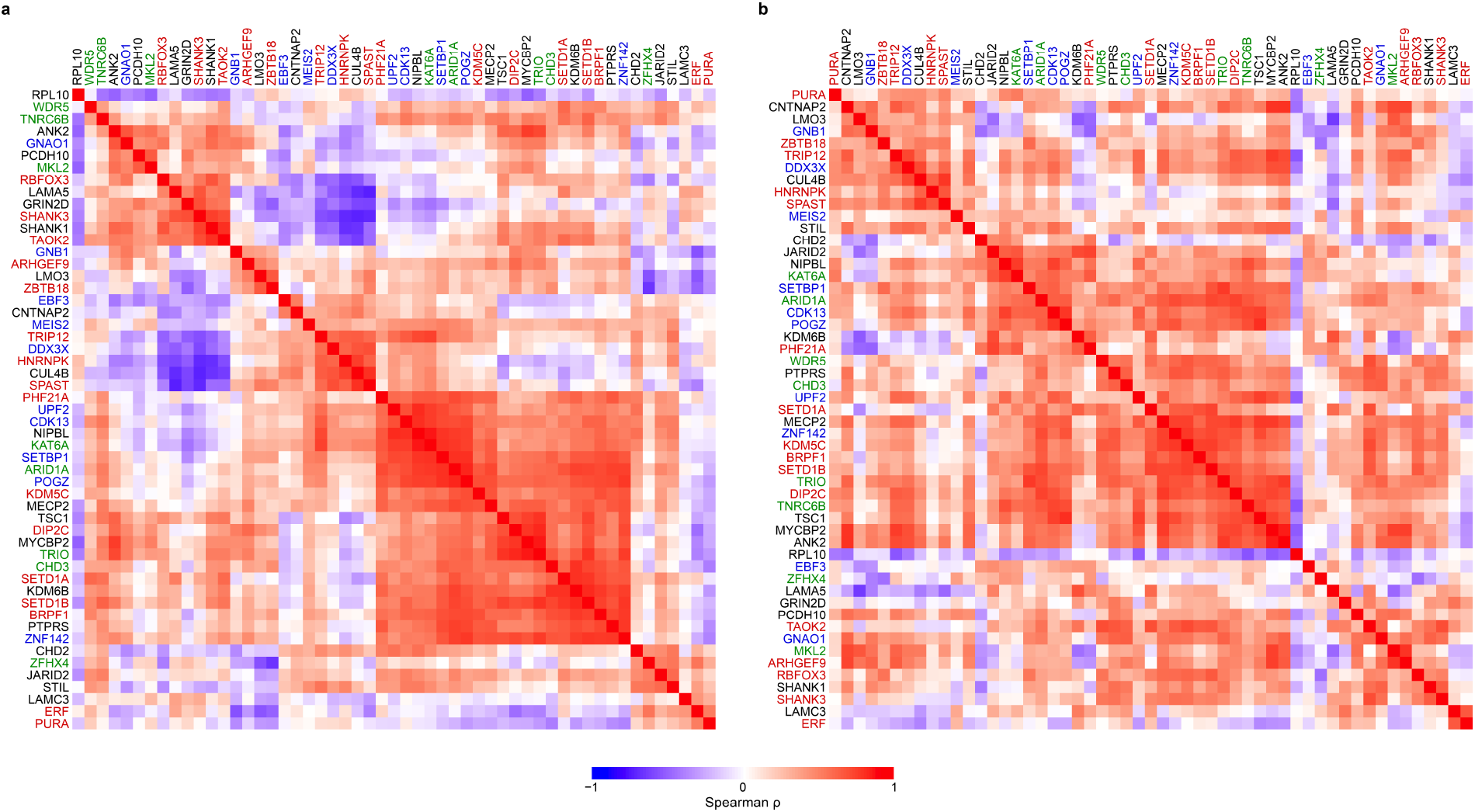
Developmental co-expression of childhood apraxia of speech candidate genes. (a, b) Spearman co-expression matrices for 53 genes in the all-region BrainSpan dataset, comprising 237 prenatal tissue samples from 20 donors (a) and 287 postnatal tissue samples from 22 donors (b). The network includes 20 research-prioritized childhood apraxia of speech (CAS) candidate genes and genes reported in previous CAS studies. Label colors indicate displayed source assignments: present study, black; Eising et al. (2019), green; Hildebrand et al. (2020), blue; and Kaspi et al. (2023), red. Gene-set memberships overlap; colors do not indicate exclusive membership. The present-study candidate *SETD1A* retains its prior-study color. The former symbol *ZNF238* is displayed as *ZBTB18*. The shared color scale represents Spearman correlation (ρ). Differences between the sampled networks are interpreted descriptively (Supplementary Tables S16–S18)

Regional BrainSpan analysis demonstrated heterogeneous developmental expression of the 20 high-confidence CAS candidate genes across cerebellar, pre-rolandic motor, primary somatosensory, and basal ganglia/frontostriatal systems. Prenatal profiles showed marked gene- and region-specific variation, with different candidates reaching relative expression peaks in distinct anatomical systems and developmental intervals. After 24 PCW, expression remained detectable across all four regional groups but generally followed more sustained, gene-specific trajectories extending through childhood and adulthood. The regional profiles therefore support a distributed neurodevelopmental model of CAS in which heterogeneous genetic disruptions affect multiple components of the speech-motor network at different developmental stages (Supplementary Tables S13–S15).

The volcano analyses further showed that most high-confidence CAS genes differed across developmental periods, with 17 detected at nominal *P* < .05 and 16 at *P* < .01. Of the 17 candidates meeting nominal P < .05, 13 showed higher expression at 8–24 PCW, and 10 showed at least 1.5-fold prenatal enrichment (Supplementary Table S12). Together, these findings indicate that genetically heterogeneous CAS candidates participate in developmentally reorganized transcriptional networks spanning brain regions central to speech-motor planning, sensorimotor integration, and neural-circuit maturation. Although overall co-expression was stronger after 25 PCW, high-confidence CAS candidates showed a pronounced early-developmental expression bias. The strongest early-developmental signals were observed for *KDM6B, LMO3, RPL10, PTPRS, CNTNAP2,* and *NIPBL*, each of which showed more than twofold higher prenatal expression. Together, these findings distinguish two complementary developmental patterns: many high-confidence CAS candidates are most highly expressed prenatally, whereas their broader transcriptional network becomes more positively coordinated during later development. This temporal reorganization supports convergence of genetically heterogeneous CAS-associated genes on regulatory and neural-circuit pathways relevant to speech-motor development. Thus, the later-development network displayed greater overall coordination, whereas many CAS candidate genes reached their highest expression during developmental periods associated with neurogenesis, chromatin regulation, neuronal differentiation, and early circuit formation.

### Functional analysis of prioritized gene candidates

While the BrainSpan heatmaps show where and when our proposed high-confidence CAS genes are naturally expressed during human brain development, they do not provide information as to why and how the co-expressed genes function together. To gain insights into the functional connections between our high-confidence genes, we conducted STRING enrichment analysis and protein-protein interaction network analysis to investigate potentially shared known developmental and neurodevelopmental biological processes. Finally, we connect our STRING analysis to the KOLF2.1J Perturb-seq atlas in order to study whether experimental repression of one gene produces downstream transcriptional responses in other genes, providing information distinct from endogenous BrainSpan expression and annotated STRING associations.

STRING Gene Ontology analysis identified enrichment of several biological processes related to nervous system development and neuronal organization among the 20 prioritized candidate genes. Nervous system development included 16 of the 20 genes (80%; FDR = 2.76 × 10⁻⁸), and neuron projection development included six (30%; FDR = 0.0254). Regulation of cell junction assembly included four genes (20%; FDR = 0.0270), while positive regulation of histone H3-K9 methylation included *JARID2* and *MECP2* (2/20, 10%; FDR = 0.0409). Percentages indicate the proportion of the 20 input genes annotated to each term; categories overlap and therefore do not sum to 100%. These findings support shared annotated biological processes within the prioritized gene set involved in early brain development, neuronal connectivity, and transcriptional regulation (Supplementary Table S26). These enrichments involved multiple CAS candidates, including *SETD1A, KDM6B, GRIN2D, NIPBL, TSC1, JARID2, CNTNAP2, CHD2, CUL4B, MECP2, RPL10,* and novel candidates *MYCBP2* and *PTPRS*. Disease-gene enrichment implicated autism spectrum disorder, epilepsy, and other brain disorders (Supplementary Table S25), while human-phenotype enrichment included autistic behavior, seizures, hyperactivity, neurodevelopmental delay, delayed speech and language development, language impairment, hypotonia, and behavioral abnormalities (Supplementary Table S27). EGF-domain enrichment involving *LAMA5, LAMC3,* and *CNTNAP2* provided additional evidence for extracellular-matrix and cell-adhesion involvement.

STRING protein–protein interaction analysis of the 20 high-confidence CAS candidate genes identified a significantly enriched network comprising 20 nodes and 63 observed edges, compared with 31 edges expected by chance (protein-protein interaction enrichment *P* = 2.61 × 10⁻⁷; Supplementary Table S23). The network had an average node degree of 6.3 and a mean local clustering coefficient of 0.681, indicating substantial functional connectivity among the prioritized genes. Gene-level connectivity statistics are provided in Supplementary Table S24. The network (Fig. 6a) suggested three broad, interconnected biological layers, starting with LAMA5 and LAMC3 located in the extracellular matrix are likely part of a Laminin-513 complex (see below) and are thus one specific unit within the network, shown as squares in Fig. 6a. The CNTNAP2 protein connects Laminin-513 to a network comprised of membrane and synaptic organization, intracellular signaling and neuronal excitatory function (Fig. 6a, triangles). Connected primarily via ANK2 and TSC1 is this signaling network to regulation of gene expression (Fig. 6a, circles). Thus, as observed previously (Raaz et al. 2026), LAMA5, the laminin-α subunit, provides the extracellular access to the neuronal cells. While we did not find the β-subunit in the present families, LAMB1 was associated with CAS in our previously analyzed patients (Raaz et al. 2026). The current cohort now identifies the previously missing link, the γ-subunit as LAMC3. Thus, we can now refine the laminin structure to be a laminin-513, instead of a laminin-511 that we had previously proposed due to lack of data on the γ-subunit identity (Fig. 6b). This is important because we have not seen any integrin receptor in any CAS-associated genes so far (Raaz et al. 2026) which can now be explained by the fact that LAMC3 is missing the critical glutamic acid present in LAMC1 and LAMC2, required for the interaction with integrin receptors (Ido et al. 2008). This structural analysis not only supports the role all three genes play in connecting neuronal cells to their extracellular environment through formation of a structural complex, but also allows visualization of the potential impact the CAS-associated mutations (see Table 1) have on this structure. Shown in Fig. 6b are the locations of the CAS mutations found in LAMA5 (p.Leu3518ProfsTer25) and in LAMC3 (T101I). For reference, our previously identified mutation in LAMA5 (R2659W) is also shown. While the R2659W likely directly interferes with core helical bundle formation (Raaz et al. 2026), the new mutations reported here likely work through interference of oligomerization patterns between EGF domains and helical bundle domains. To test this hypothesis, we deleted only 178 out of 3,695 total amino acids in LAMA5, namely those C-terminal amino acids predicted to be lost by the p.Leu3518ProfsTer25 mutation. The deletion has a dramatic effect on structure prediction of the Laminin-513 complex (see Supplementary Figure S2), which also impacts the position of the LAMC3 mutation, T101I. Thus, both mutations reported here may act through similar mechanisms despite being in different proteins and different secondary structure and domain elements.

**Table 1.**
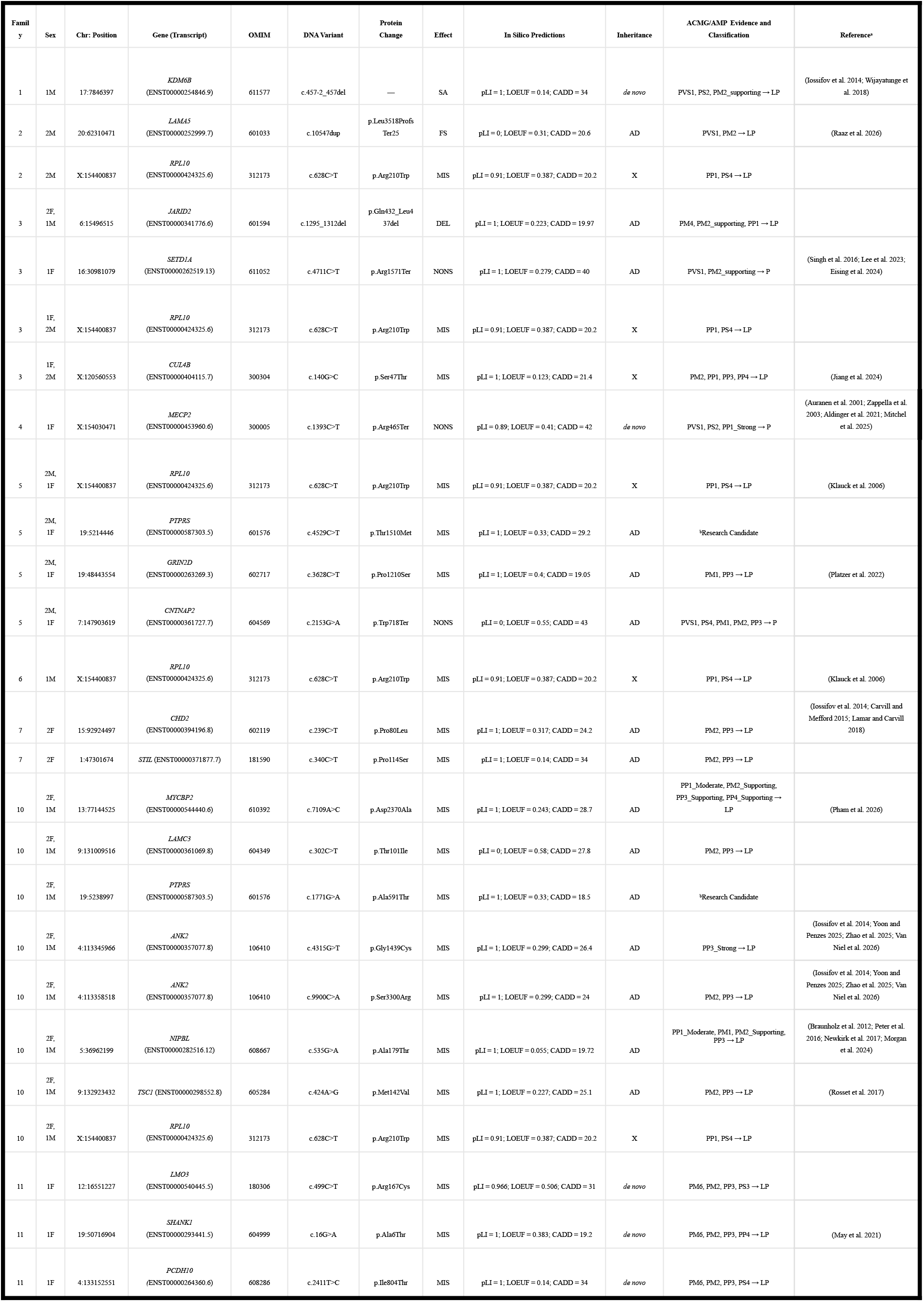
High-confidence gene variants identified in individuals with childhood apraxia of speech. OMIM, Online Mendelian Inheritance in Man; ACMG, American College of Medical Genetics; NA, not applicable; P, pathogenic; LP, likely pathogenic; VUS, variant of uncertain significance. All coordinates correspond to the *Homo sapiens* (human) genome assembly GRCh38 (hg38) from Genome Reference Consortium. In silico pathogenicity predictions: pLI and LOEUF are gene-level constraint metrics obtained from gnomAD; CADD is a variant-level deleteriousness prediction. Abbreviations for variant types follow Sequence Ontology (SO) and Human Genome Variation Society (HGVS) guidelines: splice acceptor variant (SA / splice_acceptor_variant), frameshift variant (FS / frameshift_variant), missense variant (MIS / missense_variant), in-frame deletion (DEL / inframe_deletion), and stop gained (NONS / stop_gained). Additional abbreviations and prediction thresholds: CADD (Combined Annotation Dependent Depletion), gnomAD, Genome Aggregation Database; LoF, loss of function; pLI, probability of loss-of-function intolerance; ≥ .90 indicate high intolerance to loss-of-function; LOEUF, loss-of-function observed/expected upper bound fraction; lower values indicate greater loss-of-function constraint. Abbreviations for Inheritance types: AD (Autosomal Dominant), X (X-linked). In the Sex column, the number indicates the number of individuals within the family, and the letters indicate male (M) or female (F). ᵃPreviously reported variant. ᵇResearch-candidate status denotes prioritization for further investigation.

**Table 2.** Clinical and neurodevelopmental characteristics of affected individuals with childhood apraxia of speech.

| Family | Age<sup>a</sup> at Testing | Sex | Gross motor delay | Fine motor delay | Vision impaired | Hearing loss | Seizures | Dysmorphic features, oral motor, and motor speech findings |
| --- | --- | --- | --- | --- | --- | --- | --- | --- |
| 2 | 8;3 | M | Y | Y | N | N | N | Slow alternating smile-pucker movements and slow alternation between /i/ and /u/; swallowing appeared functional. |
| 2 | 10;7 | M | Y | Y | N | N | N | Difficulty with lip protrusion; effortful alternating smile-pucker movements and /i/-/u/ productions; difficulty elevating the tongue tip to the alveolar ridge; lateral jaw movement accompanied tongue lateralization; unable to swallow water while maintaining a smile. |
| 3 | 28;9 | F | N | N | N | N | N | Difficulty with alternating smile-pucker movements and /u/-/i/ productions; lip pursing observed during swallowing; unable to maintain a smile while swallowing water. |
| 5 | 10;2, 10;5 | M | Y | N | N | N | N | Difficulty with lip protrusion; effortful alternating smile-pucker movements and /i/-/u/ productions; lateral jaw movement accompanied tongue lateralization; compensatory lip movement observed during swallowing. |
| 5 | 7;4, 7;7 | F | Y | N | N | N | N | Difficulty with alternating smile-pucker movements; production of /u/ and /i/ was comparatively easier; reduced tongue-jaw dissociation during tongue lateralization; difficulty with the smile-and-swallow task. |
| 5 | 4;3, 4;6 | M | Y | N | N | N | N | Unable to puff the cheeks; difficulty alternating between /i/ and /u/; unable to produce /f/; lateral jaw movement accompanied tongue lateralization. |
| 10 | 18;11 | F | Y | Y | N | N | N | Oral-motor performance was within normal limits, with successful completion of the smile-and-swallow task. Tonsils and uvula could not be adequately visualized during the virtual assessment. |
| 10 | 14;7 | F | Y | Y | N | N | N | N/A |
| 10 | 7;4, 7;7, 7;9 | M | Y | Y | N | N | N | Difficulty alternating between /u/ and /i/; tongue elevation was performed with the tongue outside the oral cavity; reduced tongue-jaw dissociation during tongue lateralization; difficulty with the smile-and-swallow task. |

<sup>a</sup>Age is reported as years; months. Multiple ages indicate repeated assessments conducted at different time points.

The membrane-associated component was particularly interconnected around *CNTNAP2, SHANK1, ANK2, PTPRS,* and *PCDH10* with neuronal signaling and structure, consistent with convergence on neuronal adhesion, extracellular-matrix interactions, and synaptic organization. *ANK2* and *TSC1* occupied comparatively central positions between the membrane/synaptic and regulatory components, suggesting potential bridging roles linking neuronal structure and signaling with downstream cellular programs. A dense chromatin and transcriptional-regulatory component included *NIPBL, CHD2, KDM6B, SETD1A, JARID2,* and *MECP2*. By comparison, *STIL* (*STIL* Centriolar Assembly Protein) was relatively isolated and *LMO3* showed limited connectivity, suggesting that their potential biological relevance may not be fully represented by currently annotated protein-association evidence. Separately, *STIL* ranked eighth highest out of 38,606 genes in KOLF2.1J Perturbation Cell Atlas for Cluster 22 (Supplementary Table S22). Cluster 22 dynamically tracks the pre-mitotic and mitotic fidelity network. *STIL* acts along co-clustered mitotic genes that are upregulated during cell division to manage the replication of centrioles. *STIL* is highly sensitive to mitotic perturbations; disruption of *STIL* severely impairs mitotic progression and cell survival in KOLF2.1J. *STIL* notably displayed marked depletion by day 14 (fitness *z* = −2.10).

Perturbation analysis (Fig. 6c, using the same grouping symbols as Fig. 6a) reveals additional directional regulatory relationships between prior CAS neurodevelopmental gene sets and our 20 high-confidence CAS candidate genes. KOLF2.1J perturbation atlas analysis provided an orthogonal functional-regulatory comparison between the present high-confidence CAS candidate genes and previously reported CAS/neurodevelopmental gene sets (Supplementary Table S21). Although exact gene overlap across the present cohort and the Eising, Kaspi, and Hildebrand datasets was limited, with *SETD1A* representing the principal shared gene, perturbation-based analysis identified additional directional relationships between gene sets. All 20 high-confidence CAS candidates were represented as measured expression targets, and 13 were directly perturbed (Supplementary Table S19). Directional relationships between present CAS candidates and previously reported speech and neurodevelopmental genes are provided in Supplementary Table S21. The HDBSCAN cluster-level rankings and NTC z-scores are provided in Supplementary Table S22.

Given the multilocus profile observed in Family 10, the perturbation analyses were examined for functional relationships involving *NIPBL*, *ANK2*, *LAMC3*, and the other prioritized candidates. In the KOLF2.1J hiPSC atlas, NIPBL was one of six perturbations assigned to the Cohesin-complex cluster. Within this cluster, *LMO3* had a positive mean NTC z-score of 0.52 and ranked fourth among 38,606 measured expression features, whereas *ANK2* had a negative mean NTC z-score of −0.12 and appeared among the 50 lowest-ranked features. Separately, the individual *NIPBL* perturbation yielded a positive *LMO3* response score of 0.56. These scores and rankings describe atlas-derived transcriptional responses; the rankings do not establish statistical significance, direct regulation, or the effects of the patient-specific variants. The individual-perturbation and cluster-averaged observations represent related summaries of the same atlas rather than independent replication. Biologically, the cohesin complex is a conserved chromatin-organizing protein machine that helps shape three-dimensional genome architecture by organizing DNA loops and regulating enhancer-promoter communication. Because cohesin influences chromatin structure and transcriptional control, the association of *LMO3* with the Cohesin-complex cluster suggests that *LMO3* may be responsive to perturbations affecting chromatin organization and gene-regulatory architecture. Because *LMO3* was identified as a *de novo* CAS candidate in the present cohort, this result prioritizes a testable association between *LMO3* expression and cohesin-related perturbation responses (Supplementary Table S22). The findings strengthen evidence that genetically heterogeneous CAS candidates may converge on cohesin-mediated genome organization and downstream transcriptional programs.

*LMO3* and *RPL10* reflected opposing patterns indicating that CAS-associated genes participate in distinct, context-dependent regulatory responses. *RPL10* showed negative cluster-averaged expression-response scores in several perturbation contexts, including the TFIID-complex cluster (NTC z = −0.40) and the Heparan Sulfate Proteoglycan cluster (NTC z = −0.08). These observations describe response direction and magnitude without establishing statistical significance or direct regulation (Supplementary Table S22). Conversely, *LMO3* was among the most negative features of the SIN3 complex cluster, demonstrating that its response varied across perturbation contexts. Perturbations within the Heparan Sulfate Proteoglycan cluster elicited opposing transcriptional responses among CAS candidates, with increased *LMO3* and decreased *RPL10* expression, linking these genes to extracellular-matrix, cell-adhesion, and neurodevelopmental signaling contexts.

Across the complementary analyses, the CAS candidates showed coordinated developmental expression, significant protein-association and functional enrichment, and shared perturbation-response patterns. BrainSpan co-expression included chromatin and transcriptional regulators such as *SETD1A, KDM6B, CHD2, JARID2, CUL4B, MECP2,* and *NIPBL*. STRING analysis identified enrichment involving nervous-system development, transcriptional regulation, neuronal adhesion, extracellular-matrix biology, and synaptic processes, including an EGF-domain feature involving *LAMA5, LAMC3,* and *CNTNAP2*. KOLF2.1J analysis identified additional directional and cluster-level relationships involving cohesin-associated regulation, extracellular-matrix and cell-adhesion signaling, protein homeostasis, and neurodevelopmental transcriptional responses. Together, these results identify several interconnected biological contexts for the genetically heterogeneous CAS candidates without implying that all candidates act through a single pathway (Supplementary Tables S16–S27).

Collectively, these results support convergence of genetically heterogeneous CAS candidates on interconnected pathways governing transcriptional regulation, brain development, neuronal adhesion, and synaptic function, consistent with previous evidence implicating epigenetic and transcriptional dysregulation in childhood speech disorders.

## Discussion

### Genetic architecture and inheritance

The proportion of probands with prioritized candidate variants exceeded published CAS sequencing yields, although differences in ascertainment and variant-classification criteria limit direct comparison (Morgan et al. 2024). Unlike predominantly trio-based cohorts enriched for *de novo* findings—including Kaspi et al. (2023), in which 15 of 18 high-confidence findings were *de novo*—this cohort contained predominantly inherited variation. This yield should be interpreted cautiously because the small, clinically selected cohort included several multiplex families with affected siblings and extended relatives, unlike studies focused primarily on unrelated probands or trios. The presence of multiple prioritized variants in several families suggests that oligogenic or cumulative susceptibility may contribute to CAS rather than a single causal variant in every family.

Several prioritized genes encode proteins involved in chromatin regulation or transcriptional control. Pathogenic germline variation in epigenetic writers, erasers, readers, and chromatin regulators can produce neurodevelopmental disorders in which the primary sequence variant has broader downstream effects on chromatin organization and gene expression (Fahrner and Bjornsson 2014; Bjornsson 2015). Epigenetic states encompass stable changes in gene activity or chromatin state that do not require changes in DNA sequence and may be maintained through mitotic cell divisions (Mulholland and Nishiyama 2025). This framework is particularly relevant to speech and language development. During neurodevelopment, such mechanisms contribute to the maintenance of transcriptional programs as neural progenitors proliferate and differentiate, providing a potential route through which disruption of chromatin-regulatory genes could produce persistent effects on neuronal development and function. St John et al. (2023) systematically reviewed chromatin-related neurodevelopmental disorders and identified communication impairment as a prominent feature, with chromatin writers and readers appearing to play an important role in motor-speech development. Accordingly, the relevance of methylation and chromatin biology in the present cohort does not imply a separate non-Mendelian etiology; instead, these processes provide plausible downstream mechanisms through which germline sequence variants may alter developmentally regulated transcriptional programs relevant to speech-motor circuitry. DNA methylation, histone modifications, and chromatin state were not directly measured in the participants; these relationships invite functional and epigenomic validation.

The concentration of prioritized genes involved in chromatin modification, transcriptional regulation, and higher-order chromatin organization—including *KDM6B, MECP2, SETD1A, JARID2, CHD2, CUL4B,* and *NIPBL*—is consistent with this genetic-to-regulatory framework. Previous reports implicating *NIPBL* in familial CAS (Peter et al. 2016) and *SETD1A* in independent CAS cohorts (Eising et al. 2019; Kaspi et al. 2023) further place these findings within a broader group of chromatin- and transcription-associated CAS genes. *NIPBL*, *ANK2*, and *LAMC3* link cohesin-mediated chromatin organization, neuronal membrane–cytoskeletal scaffolding, and laminin-associated extracellular-matrix and cortical-development biology, respectively, within a possible multilocus susceptibility model.

Pathogenic variants in *KDM6B* have been associated with a neurodevelopmental disorder that includes speech and language impairment (Rots et al. 2023), providing context for investigating the *de novo* variant identified in the present cohort. KDM6B is a histone demethylase that promotes transcriptional activation by removing the repressive H3K27me3 mark deposited by Polycomb repressive complex 2 (Agger et al. 2007). Experimental studies implicate KDM6B in the activation of differentiation-associated genes during neural commitment and other developmental processes (Burgold et al. 2008; Sen et al. 2008). These findings provide a rationale for investigating whether the *de novo KDM6B* variant identified here disrupts developmental gene regulation.

The established role of *MECP2* in linking DNA methylation to transcriptional repression offers a related mechanistic hypothesis, although the effect of the variant identified here remains untested (Nan et al. 1998). Candidates involved in synaptic, cytoskeletal, and extracellular-matrix biology suggest that altered gene regulation and circuit formation may represent complementary routes to susceptibility. Within this framework, *PTPRS* and *MYCBP2* warrant investigation as exploratory candidates linking neuronal adhesion and signaling to speech-related neurodevelopment. These biological relationships motivate testing of both individual variant effects and possible multilocus contributions, without establishing a shared causal mechanism.

We propose *LMO3*, *PCDH10*, *MYCBP2*, and *PTPRS* as candidate genes for CAS, with distinct backgrounds of prior evidence. *LMO3* and *PCDH10* (Wei et al. 2019) are HAR-associated and were included in the literature-derived gene list accompanying Kaspi et al. (2023), while *PTPRS* represents a previously unreported CAS candidate with potential relevance to neuronal adhesion and signaling and *MYCBP2* (Pham et al. 2026) encodes an atypical E3 ubiquitin-protein ligase that is essential for proper neural development, regulating cytoskeletal dynamics required for building central nervous system axon tracts (Table 1). The present findings thereby further expand the candidate-gene landscape.

The rarity of the prioritized variants in *LMO3*, *SHANK1*, and *PCDH10* warrants further investigation but does not establish pathogenicity or de novo origin. The previous report of *PCDH10* p.Ile804Thr in individuals with Tourette syndrome provides a point of comparison across neurodevelopmental phenotypes, although its contribution to either condition remains unresolved (Depienne et al. 2019). Establishing the relevance of these variants to CAS will require additional segregation evidence and variant-specific functional studies in developmentally relevant neural models.

#### Developmental and functional convergence

The BrainSpan expression patterns suggest that the prioritized genes may contribute to developmental processes relevant to CAS beginning prenatally (Fig. 4a, b). Expression across regions involved in motor planning, sensorimotor integration, timing, and sequencing provides a rationale for investigating distributed neural systems in speech-motor development (Fig. 4a, b; Supplementary Tables S6–S7). These observations offer developmental and anatomical context for candidate-gene prioritization, although expression alone does not establish involvement in speech-specific circuits. Because BrainSpan comprises cross-sectional bulk regional tissue, these analyses cannot resolve cell-type-specific functions or establish causal mechanisms.

**Fig. 4.**
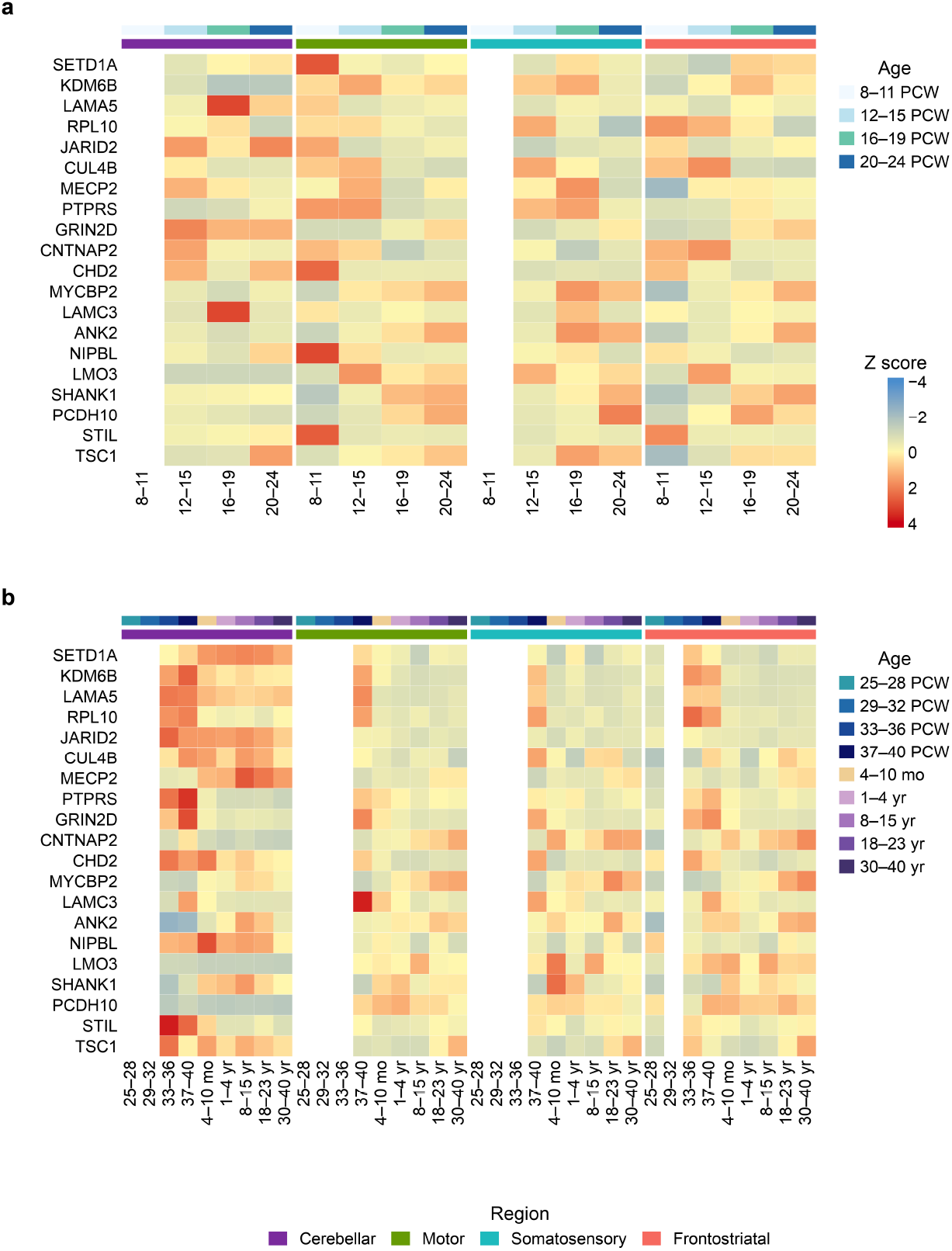
Regional developmental expression of childhood apraxia of speech candidate genes. **(a, b)** Z-score-normalized expression of 20 research-prioritized childhood apraxia of speech (CAS) candidate genes in selected BrainSpan regions at 8–24 postconceptional weeks (PCW; a) and 25 PCW through 40 years (b). Expression was averaged within region–age bins and normalized across bins for each gene; white cells indicate unavailable samples. Upper annotation strips indicate age bins, with separate age-color keys for each panel; lower strips indicate regional groups. Regions include cerebellar cortex/cerebellum (purple; CBC, CB), pre-rolandic and primary motor cortex (green; M1C, M1C–S1C), postcentral and primary somatosensory cortex (turquoise; S1C), and basal ganglia/frontostriatal regions (salmon; STR, DFC, VFC, MFC, OFC). Both panels use the shared Z-score scale and region key. Age labels are in PCW unless marked mo (months) or yr (years). The regional dataset comprised 250 tissue samples from 41 unique donors (Supplementary Tables S6–S10)

**Fig. 5.**
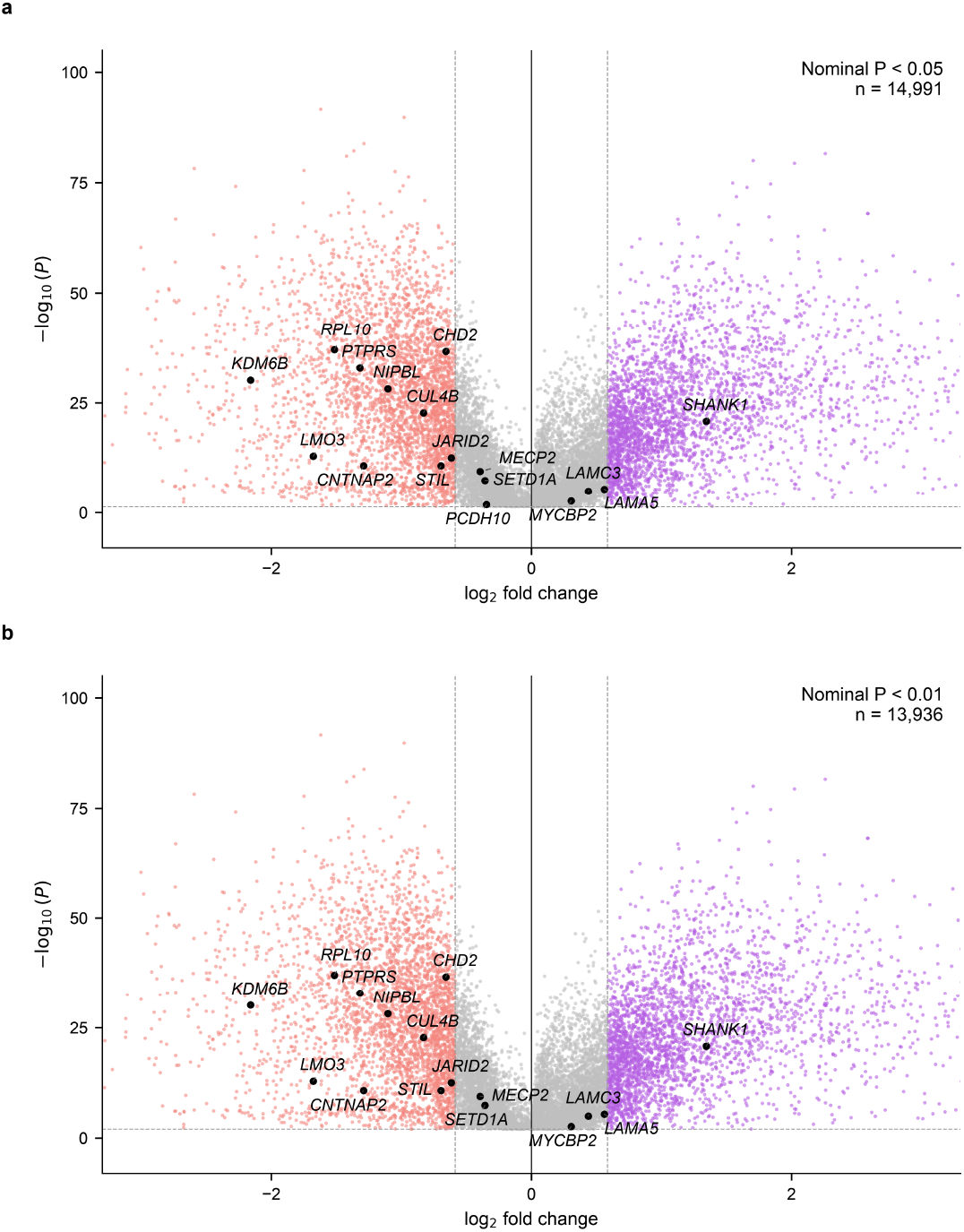
Developmental differences in BrainSpan regional gene expression. **(a, b)** Volcano plots comparing protein-coding gene expression between 8–24 postconceptional weeks (PCW; 101 tissue samples) and 25 PCW through 40 years (149 tissue samples) in the childhood apraxia of speech (CAS)-relevant regional BrainSpan dataset. Panels display features meeting nominal P < .05 (a; 14,991 features, including 17 research-prioritized CAS candidates) or P < .01 (b; 13,936 features, including 16 candidates). The x-axis shows log₂[(later mean RPKM + 1)/(early mean RPKM + 1)], and the y-axis shows −log₁₀(P) from two-sided Welch two-sample t-tests on untransformed RPKM values. Salmon indicates log₂ fold change ≤−log₂(1.5), lavender indicates log₂ fold change ≥log₂(1.5), and gray indicates smaller absolute fold changes. Black labeled points identify CAS candidates; *PCDH10* meets the P < .05 threshold only. Dashed vertical lines indicate the fold-change thresholds, and the dashed horizontal line indicates the panel-specific nominal P threshold. Fold-change thresholds determine coloring, not inclusion. These exploratory tissue-level tests do not account for repeated sampling within donors. Benjamini–Hochberg-adjusted P values across 19,130 protein-coding features are reported separately (Supplementary Tables S11a, S11b, and S12)

**Fig. 6.**
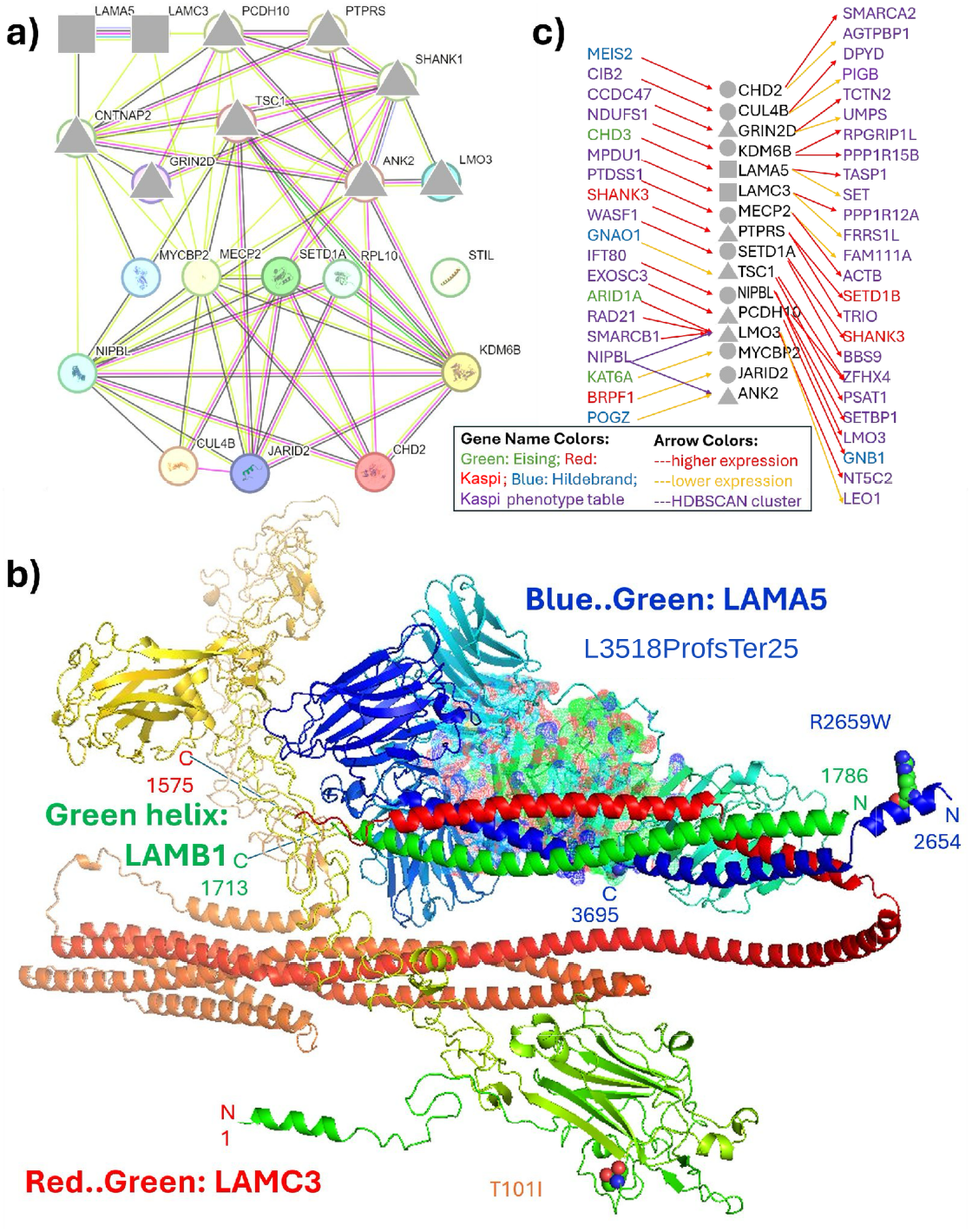
STRING network and perturbation-based functional analysis of high-confidence childhood apraxia of speech candidate genes. **(a)** STRING protein–protein association network of the 20 high-confidence CAS candidate genes. The network contained 20 nodes and 63 observed edges, compared with 31 edges expected by chance (protein–protein interaction enrichment, *P* = 2.61 × 10⁻⁷), with an average node degree of 6.3 and an average local clustering coefficient of 0.681. Enrichment implicated nervous-system and brain development, cell differentiation, autism spectrum disorder, epilepsy, seizures, neurodevelopmental delay, and delayed speech and language development (Supplementary Tables S23–S27). **(b)** Structure of the joint AlphaFold prediction of LAMA5 (C-terminal residues 2654-3695) and LAMB1 (residues 1713-1786, the same central bundle helical stretch as shown in the Laminin-511 structure with PDB 5XAU) and full-length LAMC3; observed in the AlphaFold prediction in an analogous position to chain C of 5XAU representing LAMC1 in Laminin-511, supporting its role as the γ-subunit in Laminin-513 in direct analogy to Laminin-511. The image is shown in the same orientation as in our previous paper (Figure 6d, ref. Raaz et al. 2026) but using full-length LAMC3, instead of the short helical fragment of LAMC1 from 5XAU. Highlighted in spheres are the mutations described in this paper and in Raaz et al. 2026 associated with CAS. The LAMA5 mutation reported here p.Leu3518ProfsTer25 results in a loss of the amino acid sequence until the C-terminus highlighted as dotted spheres to indicate the absence of amino acids. **(c)** Selected transcriptional-response associations from the KOLF2.1J Perturbation Cell Atlas connecting present CAS candidates with genes from previous CAS studies and the broader Kaspi phenotype-associated comparison sets (Eising et al. 2019; Hildebrand et al. 2020; Kaspi et al. 2023). The left panel contains 19 comparison-set genes; the central panel contains 13 directly perturbed CAS candidates and three additional expression targets (*MYCBP2, JARID2,* and *ANK2*); and the right panel contains 25 response genes. Red and yellow arrows indicate higher and lower expression following perturbation, respectively, and purple lines indicate HDBSCAN cluster membership. Gene-label colors indicate comparison-set membership. Central circles, squares, and triangles denote chromatin/cohesin, synaptic/adhesion/extracellular-matrix, and signaling/excitability categories, respectively. *NIPBL* was a member perturbation of the Cohesin-complex cluster, in which *LMO3* ranked fourth among positive response features (average non-targeting control [NTC] z-score = 0.52) and *ANK2* ranked 40th among negative response features (average NTC z-score = −0.12). Separately, individual *NIPBL* perturbation was associated with higher *LMO3* expression (NTC z-score = 0.56). NTC z-scores describe transcriptional responses relative to non-targeting controls and are descriptive response measures rather than *P* values or fitness measurements. Relationships were selected for illustration, with no more than two downstream targets displayed per perturbed CAS candidate. Rankings and visualization thresholds were interpreted descriptively rather than as statistical-significance criteria. Supplementary Tables S20, S21a, S21b, and S22 provide reverse-direction relationships, the complete retained downstream relationship set, the displayed downstream subset, and cluster responses, respectively.

STRING analysis provided complementary evidence of functional convergence. In particular, the involvement of laminin-related genes such as *LAMA5* and *LAMC3* suggests convergence between extracellular-matrix organization and processes involved in neuronal migration, cortical development, and circuit formation, consistent with the established neurodevelopmental relevance of *LAMC3* and additional support for *LAMA5* from Raaz et al. (2026).

#### Perturb-seq functional prioritization

Perturb-seq provides an experimental basis for developing mechanistic hypotheses beyond expression patterns and established functional associations. The relationship between *NIPBL* and *LMO3* in cohesin-associated transcriptional responses motivates testing whether altered chromatin regulation influences *LMO3*-related developmental processes. Similarly, the responses linking *PCDH10* with *GNB1* and *ZFHX4* warrant investigation of whether these genes participate in shared regulatory processes relevant to neural development. These relationships do not establish direct regulation or physical interactions: similarity in perturbation-response profiles may reflect indirect effects or broader changes in cellular state. Because KOLF2.1J cells are pluripotent, validation in differentiated neural models and comparison with patient-specific variant effects are needed to assess the relevance of these hypotheses to CAS.

### Clinical significance

This work complements previous genotype-phenotype studies not only in terms of methodology but also in contributing new insights into potential mechanisms of CAS. Since CAS disrupts children’s ability to communicate, the intense courses of interventions needed typically cannot be initiated until they are old enough to exhibit not only evidence of speech sound delays but also specific traits of CAS. A better understanding of genetic influences on CAS offers the opportunity to identify very young children at genetic risk for CAS and to offer proactive interventions that focus on precursor and earliest signals of communication. Babble Boot Camp (BBC) was developed as an early intervention and trialed in infants with classic galactosemia identified through the Recommended Uniform Screening Panel (RUSP), which enables diagnosis shortly after birth (Potter et al. 2024). If infants with genetic risk for CAS or related communication difficulties can be identified early, they may similarly become candidates for intervention before speech difficulties emerge. This opportunity is not uniform across CAS yet, however, given its genetic heterogeneity and frequent occurrence in isolation (Morgan et al. 2024). The present study contributes evidence toward this goal by showing not only genotype-phenotype associations but also interactive processes converging on the early developing brain. Genetic diagnosis may provide an opportunity to identify some children at elevated communication risk before CAS features emerge, potentially expanding the population that could benefit from genetically-informed early interventions such as BBC. In the future, clinical practice can consider an individual child’s genetic risk profile and leverage the effectiveness of earliest proactive interventions toward mitigating the predicted speech and language challenges associated with the known risk for communication challenges (Peter et al. 2023).

### Future directions

The candidacy of *MYCBP2* and *PTPRS,* as well as *PCDH10* and *LMO3,* should be evaluated through replication in independent CAS and severe speech-disorder cohorts, detailed segregation and phenotype analyses, and assessment of variant-specific effects on expression or function. Studies in neural progenitors or differentiated neuronal models could determine whether the identified variants alter cell adhesion, synaptic development, or downstream signaling. Such experiments, combined with segregation and phenotype analyses, could clarify whether each proposed novel CAS candidate gene contributes through a single-gene or multilocus mechanism.

Future studies should test the regulatory relationships identified in KOLF2.1J in developmentally and phenotypically relevant human models. Patient-specific variants in *LMO3, NIPBL, PCDH10,* and other prioritized genes could be introduced into isogenic hiPSC lines by genome editing, followed by differentiation into cortical, striatal, motor, and cerebellar neuronal lineages implicated by the BrainSpan analysis. Single-cell transcriptomic and chromatin-accessibility profiling across differentiation and maturation would determine whether the observed perturbation responses persist in neural cells, are developmentally stage-specific, and reproduce the directionality detected in the atlas.

Mechanistic experiments should include reciprocal perturbations, independent guide RNAs, dose-response and time-course analyses, and rescue of altered expression or cellular phenotypes by restoration of the perturbed gene. The proposed cohesin-related connection could be evaluated using cohesin occupancy, chromatin-accessibility, enhancer–promoter interaction, and three-dimensional genome-organization assays. The *PCDH10*-associated network could be examined through neuronal adhesion, neurite growth, synapse formation, network activity, and electrophysiological assays. Replication across independent genetic backgrounds and direct comparison of variant-specific editing with complete or partial gene repression will be essential, because CRISPRi knockdown may not reproduce the molecular consequences of individual missense, truncating, splice, or regulatory variants. Collectively, these experiments would determine whether the atlas-derived relationships represent causal regulatory mechanisms contributing to CAS or broader, context-dependent transcriptional responses.

### Limitations and validation controls

This study should be interpreted in the context of several limitations. First, although the proportion of probands with prioritized variants was high, the cohort comprised only eleven families with CAS, limiting statistical power and the generalizability of the findings. Replication in larger, independent cohorts will be necessary to confirm the candidate genes identified in this study. The KOLF2.1J results therefore provide experimentally grounded hypotheses—not causal conclusions—that now enable targeted testing of how CAS-associated variation disrupts gene regulation and neural-circuit development.

Whole-exome sequencing does not capture all forms of genomic variation, including noncoding regulatory variants, structural rearrangements, repeat expansions, or epigenetic alterations. Variant-specific functional validation was beyond the scope of this study. The proposed contributions of *LMO3*, *PCDH10*, *MYCBP2,* and *PTPRS* to CAS therefore require independent cohort replication, additional segregation evidence, and functional studies in developmentally relevant neural models. BrainSpan transcriptomic analyses also rely on cross-sectional bulk tissue and cannot establish cell-type-specific functions or causal mechanisms. Nevertheless, the integration of genomic, developmental, and systems biology evidence provides a convergent rationale for prioritizing these genes for CAS-focused investigation. Validation for newly proposed candidate genes such as *LMO3, PCDH10, MYCBP2*, and *PTPRS* will come from independent cohort replication, segregation evidence, and functional studies showing a plausible impact on neuronal development or relevance to speech.

## Conclusions

This study integrates family-based exome sequencing, segregation analysis, developmental transcriptomics, protein– protein interaction networks, and perturbation data to refine the genetic architecture of CAS. The predominance of inherited candidate variation highlights the value of studying multiplex families in investigations of CAS susceptibility. Complementary developmental and functional analyses provided biological context for these findings without establishing a single shared causal pathway. Together, the findings support marked genetic heterogeneity in CAS and raise the possibility of multilocus or oligogenic susceptibility in some families, while the magnitude of individual variants’ contributions and their influence on speech-motor development require further investigation. *PCDH10, LMO3, PTPRS*, *MYCBP2* extend this landscape toward neuronal adhesion and axon–synapse regulation but remain exploratory candidates requiring independent replication and variant-specific functional validation.

## Supporting information

Online Resource 1

Online Resource 2

## Statements and Declarations

### Ethics approval

This research was conducted in accordance with the Declaration of Helsinki and under Institutional Review Board oversight at Arizona State University (STUDY00013638, Biology of Language; and STUDY00010292, Genes, the Brain, and How We Communicate) and the University of Washington (protocol 38237).

### Consent to participate

Written parental permission was obtained for all minor participants, and written informed consent was obtained from all adult participants.

### Consent for publication

The informed-consent and parental-permission procedures included permission for the use of study data in research publications and professional educational activities.

### Data availability

The data supporting the findings of this study are included in the article and its supplementary information. Individual-level genomic and clinical data are not publicly available because of participant privacy, consent, and ethical restrictions. Additional de-identified data may be made available by the corresponding author upon reasonable request and subject to applicable institutional and ethical approvals.

### Code availability

The custom code used in this study is available from the corresponding author upon reasonable request.

### Author contributions

Tiffanie White and Marjan Chapi contributed equally to this work.

Tiffanie White: Conceptualization, Methodology, Investigation, Formal Analysis, Software, Data Curation, Visualization, Writing – Original Draft, Writing – Review & Editing.

Marjan Chapi: Conceptualization, Methodology, Investigation, Formal Analysis, Writing – Original Draft, Writing – Review & Editing.

Caitlin Raaz: Investigation, Resources, Funding Acquisition, Writing – Review & Editing.

Laurel Bruce: Investigation, Analysis, Writing – Review & Editing.

Akshit Modi: Software, Data Curation, Formal Analysis.

Valentin Dinu: Analysis, Writing – Review & Editing.

Li Liu: Methodology, Writing – Review & Editing.

Judith Klein-Seetharaman: Methodology, Formal Analysis, Visualization, Writing – Review & Editing.

Beate Peter: Funding Acquisition, Writing – Review & Editing.

All authors contributed to the interpretation of the results, critically reviewed the manuscript, approved the final version, and agree to be accountable for their respective contributions.

**Online Resource 1: Supplementary Methods and Results, including detailed phenotyping, sequencing, variant-filtering, classification, and robustness procedures, and Supplementary Figs. S1 and S2.**

**Online Resource 2: Supplementary Tables S1–S27, including participant phenotypes, prioritized variants, developmental gene-expression analyses, Perturbation Cell Atlas analyses, and functional enrichment results.**

## References

Aldinger KA, Thomson Z, Phelps IG, Haldipur P, Deng M, Timms AE, Hirano M, Santpere G, Roco C, Rosenberg AB, Lorente-Galdos B, Gulden FO, O’Day D, Overman LM, Lisgo SN, Alexandre P, Sestan N, Doherty D, Dobyns WB, Seelig G, Glass I, Millen KJ (2021) Spatial and cell type transcriptional landscape of human cerebellar development. Nature Neuroscience 24:1163–1175 10.1038/s41593-021-00872-y

Agger K, Cloos PAC, Christensen J et al (2007) UTX and JMJD3 are histone H3K27 demethylases involved in HOX gene regulation and development. Nature 449:731–734. 10.1038/nature06145

American Speech-Language-Hearing Association (n.d.) Childhood apraxia of speech. https://www.asha.org/practice-portal/clinical-topics/childhood-apraxia-of-speech/. Accessed 3 May 2026

Auranen M, Vanhala R, Vosman M, Levander M, Varilo T, Hietala M, Riikonen R, Peltonen L, Järvelä I (2001) *MECP2* gene analysis in classical Rett syndrome and in patients with Rett-like features. Neurology 56:611–617. 10.1212/WNL.56.5.611

Bennett RL, French KS, Resta RG, Austin J (2022) Practice resource-focused revision: Standardized pedigree nomenclature update centered on sex and gender inclusivity: A practice resource of the National Society of Genetic Counselors. Journal of Genetic Counseling 31:1238–1248. 10.1002/jgc4.1621

Bjornsson HT (2015) The Mendelian disorders of the epigenetic machinery. Genome Res 25:1473–1481. 10.1101/gr.190629.115

Braunholz D, Hullings M, Gil-Rodríguez MC et al (2012) Isolated *NIBPL* missense mutations that cause Cornelia de Lange syndrome alter *MAU2* interaction. Eur J Hum Genet 20:271–276. 10.1038/ejhg.2011.175

Bruininks RH, Bruininks BD (2005) Bruininks–Oseretsky Test of Motor Proficiency, 2nd edn. Pearson

Burgold T, Spreafico F, De Santa F et al (2008) The histone H3 lysine 27-specific demethylase Jmjd3 is required for neural commitment. PLoS ONE 3:e3034. 10.1371/journal.pone.0003034

Carrow-Woolfolk E (2017) Comprehensive Assessment of Spoken Language, Second Edition (CASL-2). Western Psychological Services

Carvill GL, Mefford HC (2015) *CHD2*-related neurodevelopmental disorders. In: Adam MP et al (eds) GeneReviews® University of Washington, Seattle https://www.ncbi.nlm.nih.gov/books/NBK333201/. Accessed 3 May 2026

Chan ER, Benchek P, Miller G, Brustoski K, Schaffer A, Truitt B, Tag J, Freebairn L, Lewis BA, Stein CM, Iyengar SK (2024) Importance of copy number variants in childhood apraxia of speech and other speech sound disorders. Commun Biol 7:1273. 10.1038/s42003-024-06968-y

Chenausky KV, Baas B, Stoeckel R et al (2023) Comorbidity and severity in childhood apraxia of speech: a retrospective chart review. J Speech Lang Hear Res 66:791–803. 10.1044/2022_JSLHR-22-00436

den Hoed J, Fisher SE (2020) Genetic pathways involved in human speech disorders. Curr Opin Genet Dev 65:1–7. 10.1016/j.gde.2020.05.012

Depienne C, Ciura S, Trouillard O et al (2019) Association of Rare Genetic Variants in Opioid Receptors with Tourette Syndrome. Tremor Other Hyperkinet Mov (NY) 9 10.7916/tohm.v0.693

Eising E, Carrion-Castillo A, Vino A, Strand E, Jakielski KJ, Scerri TS et al (2019) A set of regulatory genes co-expressed in embryonic human brain is implicated in disrupted speech development. Mol Psychiatry 24:1065–1078. 10.1038/s41380-018-0020-x

Eising E, Vino A, Mabie HL, Campbell TF, Shriberg LD, Fisher SE (2024) Genome sequencing of idiopathic speech delay. Hum Mutat 45:9692863. 10.1155/2024/9692863

Fahrner JA, Bjornsson HT (2014) Mendelian disorders of the epigenetic machinery: tipping the balance of chromatin states. Annu Rev Genomics Hum Genet 15:269–293. 10.1146/annurev-genom-090613-094245

Fedorenko E, Morgan A, Murray E, Cardinaux A, Mei C, Tager-Flusberg H, Fisher SE, Kanwisher N (2016) A highly penetrant form of childhood apraxia of speech due to deletion of 16p11.2. Eur J Hum Genet 24:302–306. 10.1038/ejhg.2015.149

Fletcher SG (1972) Time-by-count measurement of diadochokinetic syllable rate. Speech Hear Res 15:763–770.

Formicola D, Podda I, Dirupo E, Andreucci E, Giglio S, Cipriani P, Bombonato C, Santorelli FM, Chilosi A (2024) Expanding the molecular landscape of childhood apraxia of speech: evidence from a single-center experience. Front Neurosci 18:1396240. 10.3389/fnins.2024.1396240

Goldman R, Fristoe M (2015) Goldman–Fristoe Test of Articulation 3rd edn. Pearson.

Hildebrand MS, Jackson VE, Scerri TS, Van Reyk O, Coleman M, Braden RO et al (2020) Severe childhood speech disorder: gene discovery highlights transcriptional dysregulation. Neurology 94:e2148–e2167. 10.1212/WNL.0000000000009441

Ido H, Ito S, Taniguchi Y, Hayashi M, Sato-Nishiuchi R, Sanzen N, Hayashi Y, Futaki S, Sekiguchi K (2008) Laminin isoforms containing the gamma3 chain are unable to bind to integrins due to the absence of the glutamic acid residue conserved in the C-terminal regions of the gamma1 and gamma2 chains. J Biol Chem 283:28149– 28157. 10.1074/jbc.M803553200

Iossifov I, O’Roak BJ, Sanders SJ et al (2014) The contribution of *de novo* coding mutations to autism spectrum disorder. Nature 515:216–221. 10.1038/nature13908

Iuzzini-Seigel J (2019) Motor performance in children with childhood apraxia of speech and speech sound disorders. J Speech Lang Hear Res 62:3220–3233. 10.1044/2019_JSLHR-S-18-0380

Jiang W, Zhang J, Wang M et al (2024) The X-linked intellectual disability gene *CUL4B* is critical for memory and synaptic function. Acta Neuropathol Commun 12:188. 10.1186/s40478-024-01903-y

Kaspi A, Hildebrand MS, Jackson VE et al (2023) Genetic aetiologies for childhood speech disorder: novel pathways co-expressed during brain development. Mol Psychiatry 28:1647–1663. 10.1038/s41380-022-01764-8

Klauck SM, Felder B, Kolb-Kokocinski A et al (2006) Mutations in the ribosomal protein gene *RPL10* suggest a novel modulating disease mechanism for autism. Mol Psychiatry 11:1073–1084. 10.1038/sj.mp.4001883

Kopanos C, Tsiolkas V, Kouris A, Chapple CE, Albarca Aguilera M, Meyer R, Massouras A (2019) VarSome: the human genomic variant search engine. Bioinformatics 35:1978–1980. 10.1093/bioinformatics/bty897

Lai CSL, Fisher SE, Hurst JA, Vargha-Khadem F, Monaco AP (2001) A forkhead-domain gene is mutated in a severe speech and language disorder. Nature 413:519–523. 10.1038/35097076

Lamar KM-J, Carvill GL (2018) Chromatin remodeling proteins in epilepsy: lessons from *CHD2*-associated epilepsy. Front Mol Neurosci 11:208. 10.3389/fnmol.2018.00208

Lee S, Menzies L, Hay E et al (2023) Epigenotype–genotype–phenotype correlations in *SETD1A* and *SETD2* chromatin disorders. Hum Mol Genet 32:3123–3134. 10.1093/hmg/ddad079

Manto M et al (2012) Consensus paper: roles of the cerebellum in motor control—the diversity of ideas on cerebellar involvement in movement. Cerebellum 11:457–487. 10.1007/s12311-011-0331-9

Mariën P, Ackermann H, Adamaszek M et al (2014) Consensus paper: Language and the cerebellum: an ongoing enigma. Cerebellum 13:386–410. 10.1007/s12311-013-0540-5

May HJ, Jeong J, Revah-Politi A et al (2021) Truncating variants in the *SHANK1* gene are associated with a spectrum of neurodevelopmental disorders. Genet Med 23:1912–1921. 10.1038/s41436-021-01222-w

Miller GJ, Lewis BA, Benchek P et al (2019) Reading outcomes for individuals with histories of suspected childhood apraxia of speech. Am J Speech Lang Pathol 28:1432–1447.

Mitchel MW, Oetjens M, Berry ASF et al (2025) Monogenic disorders associated with motor speech phenotypes in children and adolescents undergoing clinical exome sequencing. Genet Med 27:101374. 10.1016/j.gim.2025.101374

Morgan AT, Amor DJ, St John MD, Scheffer IE, Hildebrand MS (2024) Genetic architecture of childhood speech disorder: a review. Mol Psychiatry 29:1281–1292. 10.1038/s41380-024-02409-8

Morgan AT, Murray E, Liégeois FJ (2018) Interventions for childhood apraxia of speech. Cochrane Database Syst Rev 5:CD006278. 10.1002/14651858.CD006278.pub3

Mulholland CB, Nishiyama A (2025) Molecular mechanisms of maintenance DNA methylation. Genes Genet Syst 100:25–00073. 10.1266/ggs.25-00073

Nan X, Ng HH, Johnson CA et al (1998) Transcriptional repression by the methyl-CpG-binding protein MeCP2 involves a histone deacetylase complex. Nature 393:386–389. 10.1038/30764

National Center for Biotechnology Information (n.d.) ClinVar. https://www.ncbi.nlm.nih.gov/clinvar/. Accessed 3 May 2026

Newkirk DA, Chen YY, Chien R et al (2017) The effect of Nipped-B-like (*NIPBL*) haploinsufficiency on genome-wide cohesin binding and target gene expression: modeling Cornelia de Lange syndrome. Clin Epigenetics 9:89. 10.1186/s13148-017-0391-x

Nourreddine S, Doctor Y, Dailamy A et al (2026) A genome-scale CRISPRi perturbation atlas of human induced pluripotent stem cells. Nat Biotechnol 10.1038/s41587-026-03199-w

Pais L, Snow H, Weisburd B, Zhang S, Baxter S, DiTroia S, O’Heir E, England E, Chao K, Lemire G, Osei-Owusu I, VanNoy G, Wilson M, Nguyen K, Arachchi H, Phu W, Solomson M, Mano S, O’Leary M, Lovgren A, Babb L, Austin-Tse C, Rehm H, MacArthur D, O’Donnell-Luria A (2022) seqr: a web-based analysis and collaboration tool for rare disease genomics. Human Mutation https://www.ncbi.nlm.nih.gov/pmc/articles/PMC9903206

Parenti I, Diab F, Ruiz Gil S et al (2020) *MAU2* and *NIPBL* variants impair the heterodimerization of the cohesin loader subunits and cause Cornelia de Lange syndrome. Cell Rep 31:107647. 10.1016/j.celrep.2020.107647

Peter B, Button L, Stoel-Gammon C, Chapman K, Raskind WH (2013) Deficits in sequential processing manifest in motor and linguistic tasks in a multigenerational family with childhood apraxia of speech. Clin Linguist Phon 27:163–191. 10.3109/02699206.2012.736011

Peter B, Wijsman EM, Nato AQ Jr et al (2016) Genetic candidate variants in two multigenerational families with childhood apraxia of speech. PLoS One 11:e0153864. 10.1371/journal.pone.0153864

Peter B, Lancaster H, Vose C, Middleton K, Stoel-Gammon C (2018) Sequential processing deficit as a shared persisting biomarker in dyslexia and childhood apraxia of speech. Clin Linguist Phon 32:316–346. 10.1080/02699206.2017.1375560

Peter B, Bruce L, Raaz C, Williams E, Pfeiffer A, Rogalsky C (2021) Comparing global motor characteristics in children and adults with childhood apraxia of speech to a cerebellar stroke patient: evidence for the cerebellar hypothesis in a developmental motor speech disorder. Clin Linguist Phon 35:368–392. 10.1080/02699206.2020.1861103

Peter B, Bruce L, Finestack L et al (2023) Precision medicine as a new frontier in speech-language pathology: how applying insights from behavior genomics can improve outcomes in communication disorders. Am J Speech Lang Pathol 32:1397–1412. 10.1044/2023_AJSLP-22-00205

Potter NL, VanDam M, Bruce L et al (2024) Virtual post-intervention speech and language assessment of toddler and preschool participants in Babble Boot Camp. J Speech Lang Hear Res 67:3327–3339. 10.1044/2023_JSLHR-22-00687

Pham A, Harmon J, Thibodaux LK, Flashman LA, Curtin M (2026) First report of an inherited *MYCBP2* neurodevelopmental disorder: review of proband and parent presentation. Neurogenetics 27:8. 10.1007/s10048-025-00854-3

Platzer K, Krey I, Lemke JR (2022) *GRIN2D*-related developmental and epileptic encephalopathy. In: Adam MP, Feldman J, Mirzaa GM et al (eds) GeneReviews®. University of Washington, Seattle. Initial posting July 28, 2022 https://www.ncbi.nlm.nih.gov/books/NBK582335/

Raaz C, Bruce L, Ganapathiraju M, Klein-Seetharaman J, Liu L, Dinu V, Chapi M, Kim E, Kim Y, White T, Peter B (2026) Three Unrelated Children With Childhood Apraxia of Speech: Exome Sequencing and Functional Gene Analysis Imply a Role of Laminin-511 in Early Neurodevelopment. Case Reports in Genetics 9927839, 21 pages, 10.1155/crig/9927839

Reynolds CR, Kamphaus RW (2015) Reynolds Intellectual Assessment Scales, 2nd edn. PAR

Richards S, Aziz N, Bale S et al (2015) Standards and guidelines for the interpretation of sequence variants. Genet Med 17:405–424. 10.1038/gim.2015.30

Rosset C, Netto CBO, Ashton-Prolla P (2017) *TSC1* and *TSC2* gene mutations and their implications for treatment in Tuberous Sclerosis Complex: a review. Genet Mol Biol 40:69–79. 10.1590/1678-4685-GMB-2015-0321

Rots D, Jakub TE, Keung C et al (2023) The clinical and molecular spectrum of the KDM6B-related neurodevelopmental disorder. Am J Hum Genet 110:963–978. 10.1016/j.ajhg.2023.04.008

Sen GL, Webster DE, Barragan DI, Chang HY, Khavari PA (2008) Control of differentiation in a self-renewing mammalian tissue by the histone demethylase JMJD3. Genes Dev 22:1865–1870. 10.1101/gad.1673508

Shriberg LD, Kwiatkowski J, Mabie HL (2019) Estimates of the prevalence of motor speech disorders in children with idiopathic speech delay. Clin Linguist Phon 33:679–706. 10.1080/02699206.2019.1595731

Silk M, Petrovski S, Ascher DB (2019) MTR-Viewer: identifying regions within genes under purifying selection. Nucleic Acids Res 47(W1):W121–W126. 10.1093/nar/gkz457

Singh T, Kurki MI, Curtis D et al (2016) Rare loss-of-function variants in *SETD1A* are associated with schizophrenia and developmental disorders. Nat Neurosci 19:571–577. 10.1038/nn.4267

St John M, Tripathi T, Morgan AT, Amor DJ (2023) To speak may draw on epigenetic writing and reading: Unravelling the complexity of speech and language outcomes across chromatin-related neurodevelopmental disorders. Neuroscience and biobehavioral reviews 152:105293. 10.1016/j.neubiorev.2023.105293

Takahashi H, Takahashi K, Liu FC (2009) FOXP genes, neural development, speech and language disorders. In K. Maiese (ed), Forkhead transcription factors: Vital elements in biology and medicine (Madame Curie Bioscience Database). Landes Bioscience. https://www.ncbi.nlm.nih.gov/books/NBK7023/

Todorović S, Anton JL, Sein J, Nazarian B, Chanoine V, Rauchbauer B, Kotz SA, Runnqvist E (2024) Cortico-Cerebellar Monitoring of Speech Sequence Production. Neurobiology of language (Cambridge, Mass.), 5:701–721. 10.1162/nol_a_00113

Torgesen JK, Wagner RK, Rashotte CA (2012) Test of Word Reading Efficiency, 2nd edn. Pro-Ed

Van Niel H, Lauretta M, Baker E et al (2026) Childhood motor speech disorders: who to prioritise for genetic testing. Eur J Hum Genet 10.1038/s41431-025-01993-9

Wei Y, de Lange SC, Scholtens LH et al (2019) Genetic mapping and evolutionary analysis of human-expanded cognitive networks. Nat Commun 10:4839 10.1038/s41467-019-12764-8

Wijayatunge R, Liu F, Shpargel KB, Wayne NJ, Chan U, Boua JV, Magnuson T, West AE (2018) The histone demethylase *KDM6B* regulates a mature gene expression program in differentiating cerebellar granule neurons. Mol Cell Neurosci 87:4–17. 10.1016/j.mcn.2017.11.005

Wolf M, Denckla MB (2005) Rapid Automatized Naming and Rapid Alternating Stimulus Tests (RAN/RAS). Pro-Ed.

Worthey EA, Raca G, Laffin JJ et al (2013) Whole-exome sequencing supports genetic heterogeneity in childhood apraxia of speech. J Neurodev Disord 5:29. 10.1186/1866-1955-5-29

Yoon S, Penzes P (2025) Roles of *ANK2*/ankyrin-B in neurodevelopmental disorders: Isoform functions and implications for autism spectrum disorder and epilepsy. Curr Opin Neurobiol 90:102938. 10.1016/j.conb.2024.102938

Zappella M, Meloni I, Longo I et al (2003) Study of *MECP2* gene in Rett syndrome variants and autistic girls. Am J Med Genet B Neuropsychiatr Genet 119B:102–107. 10.1002/ajmg.b.10070

Zhao L, Qiao ZD, Jia YX, Fu JX, Li TX, Jia KR, Zhao H, Bao JP, Yang XF, Pan H, Yang GL (2025) A Mutation in the *ANK2* Gene Causing ASD and a Review of the Literature. Mol Genet Genomic Med 13:e70083. 10.1002/mgg3.70083

