## Supplementary material for "Developmental Convergence Across Diverse Genetic Architectures in Childhood Apraxia of Speech": Online Resource 1

**Online Resource 1: Supplementary Methods, Results, and Figures S1 and S2**

**Article title: Developmental Convergence Across Diverse Genetic Architectures in Childhood Apraxia of Speech**
**Journal:** *Behavior Genetics*
**Authors:**

[†] **These authors contributed equally to this work.**

1 Krieger School of Arts and Sciences, The Johns Hopkins University, Baltimore, MD, USA

2 College of Health Solutions, Arizona State University, Phoenix, AZ, USA

3 Department of Communication Sciences and Disorders, University of Northern Colorado, Greeley, CO, USA

**Online Resource 1: Supplementary Methods, Results, and Figures S1 and S2**

### Supplementary Methods

#### S1 Whole-exome sequencing and variant annotation

Exome sequencing was performed using hg38-aligned data, and candidate variants were evaluated in the context of inheritance pattern, phenotypic fit, and segregation within each family. The resulting candidate gene list was refined through iterative review to prioritize the most plausible disease-associated loci for downstream interpretation. The whole-exome sequencing, variant-filtering, prioritization, and classification workflow is shown in Supplementary Fig. S1.

#### Genomic data generation

Saliva and blood samples were collected from participants using standardized collection kits. DNA extraction and whole-exome sequencing were performed at the University of Washington. Genomic DNA was extracted either from whole blood or saliva samples using standard protocols. Whole-exome sequencing (WES) was performed at the University of Washington Center for Rare Disease Research (UW-CRDR). DNA samples underwent standardized quality control procedures, including quantification, sex verification, and sample identity confirmation prior to library preparation. All samples met quality control requirements and were included in downstream analyses.

Exome libraries were prepared and enriched using the Twist Human Exome capture platform (Twist Bioscience; 36.5-Mb target region). Library concentration was assessed through fluorometric assay and molecular weight distributions were verified on the Agilent Bioanalyzer. Sequencing was performed using a 2 × 150-bp paired-end configuration. The mean library insert length was 180 ± 15 bp.

Massively parallel sequencing-by-synthesis with fluorescently labeled, reversibly terminating nucleotides was performed on the NovaSeq sequencer, and variant detection and genotyping were performed using Genome Analysis Toolkit (GATK) HaplotypeCaller v4.2.0.0. The sequencing pipeline combines Illumina software and other industry-standard software packages, including GATK, Picard, BWA, SAMtools, and in-house custom scripts. Mean on-target coverage was 35–40×, with more than 90% of the targeted exome covered at >10× and more than 80% covered at >20×. Sequence reads were aligned to the GRCh38/hg38 human reference genome and variants were called using GATK. Variants were annotated using SeattleSeq and filtered in seqr (Pais et al. 2022) using population frequency, predicted pathogenicity, inheritance, and phenotype-based criteria. Sequencing platform, read length, alignment software, and coverage metrics were provided by the University of Washington Center for Rare Disease Research. Seqr is a public online software platform that can be used to identify disease-causing genetic variants (seqr URL: https://seqr.gs.washington.edu).

### S2 Variant filtering and prioritization

Variant filtering and prioritization were performed using the seqr platform (Pais et al. 2022). Initial filtering was restricted to PASS variants meeting predefined call quality and rarity thresholds, including a Combined Annotation Dependent Depletion (CADD) score ≥15 and population allele frequency generally ≤ .01 (1%), with rarity consistent with the expected inheritance model. Candidate variants were subsequently assessed for segregation within each family. All prioritized variants underwent additional quality control review. Only variants with a minimum read depth >15× and genotype quality (GQ) score >20 were retained. Manual inspection of sequencing reads was performed using the Integrative Genomics Viewer (IGV) to confirm variant support, evaluate allelic balance, and exclude potential sequencing or alignment artifacts. Variants passing all filtering, segregation, and quality-control criteria were retained for downstream interpretation and classification. Variant pathogenicity classification was distinguished from research prioritization. ACMG/AMP classifications were interpreted in the context of the relevant established gene–disease association and inheritance mechanism (Richards et al. 2015). For exploratory candidates lacking an established association with the phenotype under investigation, sequence, population-frequency, computational, and segregation evidence was used to support research prioritization without establishing pathogenicity for CAS. In particular, the *PTPRS* variants are presented as research candidates whose clinical significance needs further validation.

### S3 Variant classification and reporting framework

Candidate genes identified in the initial analysis were re-filtered according to inheritance pattern and prioritization criteria to generate a refined list of the most promising loci. Gene-level annotations were consolidated in a summary table that included functional grouping, biological process classification, family membership, sex, affected status, proband relationship, and maternal and paternal ancestry. For shortlisting candidate variants among the identified loss-of-function (LoF) and predicted-damaging missense variants, literature-derived gene sets informed candidate prioritization and comparative annotation but were not used as an exclusionary filter; variants outside these sets remained eligible under the study's prioritization criteria (Eising et al. 2019; Hildebrand et al. 2020; Kaspi et al. 2023; St John et al. 2023; Morgan et al. 2024).

#### Initial variant review

During initial review, missense variants were evaluated using PolyPhen-2, SIFT, CADD, and missense-tolerance information. A CADD Phred score ≥15 was used as the preliminary CADD threshold. Variants passing this initial review were evaluated further using inheritance, segregation, sequencing quality, phenotypic concordance, and ACMG/AMP evidence.

#### Final variant prioritization and classification

For final prioritization, predicted-damaging missense variants were required to meet at least three of four criteria: (1) a PolyPhen-2 prediction of "probably damaging" or "possibly damaging"; (2) a SIFT prediction of "deleterious" or “deleterious (low confidence)”; (3) a CADD Phred score ≥20; or (4) a missense tolerance ratio (MTR) <1 with FDR < .05, indicating regional depletion of missense variation (Silk et al. 2019). CADD evidence did not independently determine variant confidence or pathogenicity. Predicted-damaging missense variants meeting these criteria are reported in Supplementary Table S4 (Online Resource 2). Variants were assessed according to American College of Medical Genetics and Genomics/Association for Molecular Pathology (ACMG/AMP) guidelines (Richards et al. 2015). Loss-of-function variants were evaluated separately from missense variants. High-confidence variants comprised loss-of-function and predicted-damaging missense variants classified as pathogenic (class 5) or likely pathogenic (class 4) according to ACMG/AMP guidelines, with phenotypic concordance between the gene-associated disorder and the proband. *PTPRS* with 2 variants were not formally classified as pathogenic or likely pathogenic was prioritized for further research. Low-confidence variants included variants of uncertain significance (class 3) or variants lacking phenotypic or functional support.

**S4 BrainSpan developmental co-expression and network analysis**

Spearman co-expression matrices were calculated for 53 genes using 237 prenatal tissue samples from 20 donors and 287 postnatal tissue samples from 22 donors. Prenatal samples were identified by ages expressed in postconceptional weeks; this group therefore included prenatal ages above 24 PCW. Network summaries were calculated from the 1,378 unique off-diagonal gene pairs. Undirected network edges were defined by $|\rho| \geq 0.50$, with self-correlations excluded, and connectivity was defined as the number of qualifying edges per gene. The correlation threshold defined network inclusion and was not interpreted as a statistical-significance threshold.

The primary developmental contrast was the postnatal-minus-prenatal difference in median absolute Spearman correlation. Uncertainty was estimated using 10,000 donor-cluster bootstrap replicates. Donors were sampled with replacement within each developmental group, with all tissue samples from each selected donor retained, and the gene ranks, correlation matrices, and network-level contrast were recalculated in every replicate. Approximate 95% percentile bootstrap confidence intervals were obtained from the 2.5th and 97.5th percentiles of the bootstrap contrast distribution.

Two sensitivity analyses were conducted using the 16 brain regions represented in both developmental groups. The first repeated the Spearman analysis after restriction to these shared regions. The second ranked expression within each developmental group, removed each gene’s regional mean rank, and calculated correlations between the resulting residuals. Regional means and ranks were recalculated within each bootstrap replicate.

The original sample-count-matched analysis was retained as a descriptive sensitivity analysis. In each of 1,000 iterations, 237 of the 287 postnatal tissue samples were randomly selected without replacement, and the median absolute correlation was recalculated. This analysis assessed sensitivity to unequal tissue-sample counts but did not account for repeated tissue sampling within donors or dependence among gene-pair correlations. The previous Wilcoxon comparison of gene-pair correlations was not used for statistical inference.

**Exploratory developmental expression comparisons and volcano plots**

Developmental expression comparisons used the selected BrainSpan regional dataset, comprising 101 tissue samples at 8–24 postconceptional weeks (PCW; Group A) and 149 tissue samples from 25 PCW through 40 years (Group B). For each of 19,130 protein-coding expression features, a two-sided Welch two-sample t-test compared untransformed reads per kilobase per million mapped reads (RPKM) values between groups without assuming equal variances. P values were adjusted across the 19,130 features using the Benjamini–Hochberg procedure. Supplementary Tables S11a and S11b retain features meeting nominal, unadjusted $P < .05$ and $P < .01$, respectively, and report adjusted P values separately.

Pseudocount-adjusted $\log_2$ fold changes were calculated as $\log_2$[(mean Group B expression + 1)/(mean Group A expression + 1)]. Volcano plots display these values against $-\log_{10}$ of the nominal P value. Directional coloring uses thresholds of $\pm\log_2(1.5)$; features below this magnitude threshold remain displayed when they meet the nominal P-value criterion. Prioritized CAS candidates are highlighted separately.

Multiple tissue samples originated from individual donors, and these gene-wise tests did not model within-donor dependence. The results are therefore interpreted as exploratory tissue-level comparisons. The donor-cluster bootstrap used for the separate co-expression network analysis does not adjust these gene-wise expression tests.

**S5 KOLF2.1J relationship extraction and visualization**

The KOLF2.1J atlas was examined in two directions: transcriptional responses of CAS candidates to perturbation of comparison-set genes, and responses of comparison-set genes to perturbation of CAS candidates. Comparison sets comprised the 2,147-gene phenotype-associated list and the published CAS candidate sets from Eising et al., Hildebrand et al., and Kaspi et al. (Supplementary Tables S2 and S18). Gene-set memberships could overlap.

For the CAS-to-comparison-gene analysis, the 13 CAS candidates represented as CRISPRi perturbations were queried. Comparison-set targets appearing among the atlas's 50 highest or 50 lowest expression-response scores were retained. Additional comparison-set targets with selected-matrix $|\text{NTC } z| \geq 0.25$ were included when the pair was not already represented in the top-50 results. Self-responses were excluded. This procedure yielded 533 perturbation–target relationships: 124 top-50 pairs and 409 additional matrix-only pairs.

Figure 4b was constructed as an exploratory visualization of selected relationships. To maintain readability and prevent individual perturbations from dominating the display, no more than two downstream targets were shown for each perturbed CAS candidate, yielding 26 displayed relationships. Selection considered response magnitude or rank and representation of previously reported CAS candidates; it was not a uniform selection of the two largest absolute scores. Additional measured-only CAS candidates were identified separately in the central panel. Gene-label colors indicate the source gene-set category used for display, with overlapping memberships retained in the supporting tables.

Individual-perturbation responses were distinguished from HDBSCAN cluster-averaged responses. Top-50 membership and the selected-matrix threshold were descriptive inclusion criteria, not statistical-significance thresholds. The figure is an illustrative subset and was not used to estimate network density, enrichment, or the prevalence of regulatory relationships.

### Supplementary Results

#### Variant analysis summary

Among high-confidence variants, the mean allele frequency was 0.00527%. High-confidence variants exhibited strong sequencing support, with a mean read depth of 66.36× (range: 18–141×), a mean allelic balance of 0.457, and uniformly high genotype quality scores with a mean genotype quality score of 99.

#### Donor-aware analysis of prenatal and postnatal co-expression networks

The BrainSpan co-expression analysis included 237 prenatal tissue samples from 20 donors and 287 postnatal tissue samples from 22 donors (Fig. 3a, b; Supplementary Table S16). Across the 1,378 unique gene pairs in the 53-gene network, median absolute Spearman correlation was 0.2882 in the prenatal network and 0.3381 in the postnatal network, an observed difference of 0.0499. A donor-cluster bootstrap with 10,000 replicates yielded an approximate 95% confidence interval of −0.0599 to 0.1168. Sensitivity analyses restricted to brain regions represented in both developmental groups and adjusted for regional mean ranks also yielded intervals that included zero. In a separate sample-count sensitivity analysis, 1,000 random subsets of 237 postnatal tissue samples produced a mean median absolute correlation of 0.337 (range, 0.315–0.358), and all iterations exceeded the prenatal value of 0.2882. This indicates that the observed pattern persisted after equalizing the number of tissue samples, but the analysis does not address repeated sampling within donors or dependence among gene-pair correlations. Thus, although the sampled postnatal network had a higher median absolute correlation, the donor-aware analyses did not provide clear evidence of a population-level developmental increase. Network edge counts and connectivity rankings are presented descriptively.

Using $|\rho| \geq 0.50$, the complete later-development network contained 387 edges, corresponding to a density of 28.1%. Mean connectivity was 14.6 edges per gene (median, 15; SD, 7.8; range, 0–29). *MECP2* and *ANK2* had the highest connectivity (29 each), followed by *TRIO* (28), *MYCBP2* and *DIP2C* (26 each), *ARID1A* and *SETD1B* (24 each), *ZNF142* (23), *SETBP1* and *KDM5C* (22 each), *TSC1* (21), *PTPRS*, *TNRC6B*, *CDK1*3, *GNAO1*, and *POGZ* (20 each), *CNTNAP2* and *STIL* (19 each), and *DDX3X* and *TAOK2* (18 each). These values describe the observed postnatal network; the connectivity rankings and threshold-defined edges were not tested for gene-specific statistical significance. The CAS-relevant regional subset shown in Fig. 4a, b included 250 tissue samples: 101 from 8–24 PCW and 149 from 25 PCW to 40 years.

Gene symbols were standardized according to current HUGO Gene Nomenclature Committee (HGNC) recommendations. Accordingly, *ZNF238* was reported as *ZBTB18*, its current HGNC-approved symbol; the former symbol is provided at first mention where necessary.

**Supplementary Fig. S1 Whole-exome sequencing, variant-filtering, prioritization, and classification workflow**

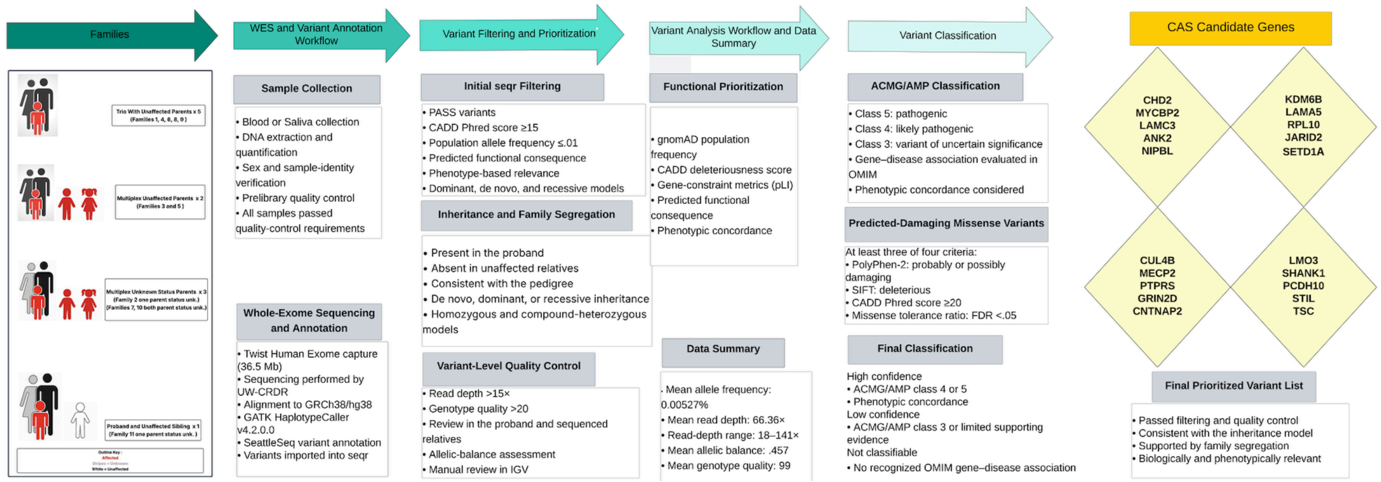


The workflow summarizes family structures, sequencing and annotation, variant filtering and prioritization, family-based analysis, ACMG/AMP classification, and candidate-gene selection. Abbreviations: ACMG/AMP, American College of Medical Genetics and Genomics/Association for Molecular Pathology; CADD, Combined Annotation Dependent Depletion; CAS, childhood apraxia of speech; FDR, false discovery rate; gnomAD, Genome Aggregation Database; GQ, genotype quality; GRCh38, Genome Reference Consortium Human Build 38; IGV, Integrative Genomics Viewer; OMIM, Online Mendelian Inheritance in Man; PASS, passed all applied variant-quality filters; pLI, probability of loss-of-function intolerance; seqr, web-based genomic analysis platform; UW-CRDR, University of Washington Center for Rare Disease Research; WES, whole-exome sequencing.

Variant filtering and prioritization were performed using seqr (https://seqr.gs.washington.edu; Pais et al. 2022), and variant classification was based on ACMG/AMP guidelines (Richards et al. 2015). The workflow diagram was created using Lucidchart (Lucid Software Inc., South Jordan, UT, USA)

**Supplementary Fig. S2 Structural modeling of LAMA5 C-terminal deletion in a candidate laminin-513 assembly**

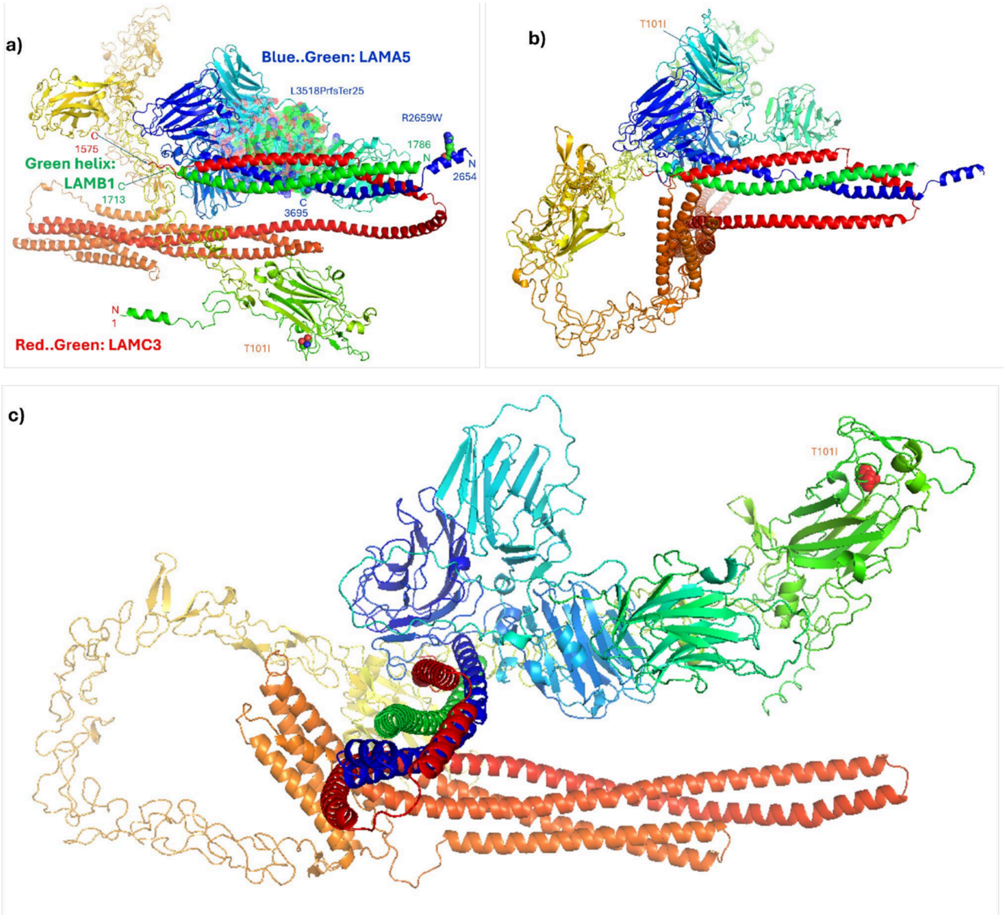


(a) Reference AlphaFold model comprising LAMA5 residues 2654–3695, LAMB1 residues 1713–1786, and full-length LAMC3. Highlighted sites correspond to the reported protein variants; dotted spheres identify the native LAMA5 C-terminal region selected for deletion. (b) Predicted assembly following removal of the final 178 residues of LAMA5, illustrating differences in the modeled arrangement of the protein chains. (c) Additional structural view highlighting the site corresponding to the LAMC3 p.Thr101Ile variant. LAMA5 is displayed in blue-to-green colors, LAMB1 as a green helix, and LAMC3 in red-to-green colors. N and C indicate protein termini. The deletion model explores loss of the native LAMA5 C-terminal segment and should be interpreted as a truncation surrogate rather than a complete reconstruction of the frameshift allele. Differences between predicted structures are hypothesis-generating and do not establish altered complex assembly or a shared pathogenic mechanism

### Supplementary Tables S1–S27 are provided in Online Resource 2.
